# Dabrafenib alters MDSC differentiation and function by activation of GCN2

**DOI:** 10.1101/2023.08.09.552588

**Authors:** M. Teresa Ciudad, Rene Quevedo, Sara Lamorte, Robbie Jin, Nadine Nzirorera, Marianne Koritzinsky, Tracy L. McGaha

## Abstract

The effect of targeted therapeutics on anti-cancer immune responses is poorly understood. The BRAF inhibitor dabrafenib has been reported to activate the integrated stress response (ISR) kinase GCN2, and the therapeutic effect has been partially attributed to GCN2 activation. Since ISR signaling is a key component of myeloid-derived suppressor cell (MDSC) development and function, we measured the effect of dabrafenib on MDSC differentiation and suppressive activity. Our data showed that dabrafenib attenuated MDSC ability to suppress T cell activity, which was associated with a GCN2-dependent block of the transition from monocytic progenitor to polymorphonuclear (PMN)-MDSCs and proliferative arrest resulting in PMN-MDSC loss. Transcriptional profiling revealed that dabrafenib-driven GCN2 activation altered metabolic features in MDSCs enhancing oxidative respiration, and attenuated transcriptional programs required for PMN development. Moreover, we observed a broad downregulation of transcriptional networks associated with PMN developmental pathways, and increased activity of transcriptional regulons driven by *Atf5*, *Mafg*, and *Zbtb7a*. This transcriptional program alteration underlies the basis for PMN-MDSC developmental arrest, skewing immature MDSC development towards monocytic lineage cells. *In vivo*, we observed a pronounced reduction in PMN-MDSCs in dabrafenib-treated tumor-bearing mice suggesting that dabrafenib impacts MDSC populations systemically and locally, in the tumor immune infiltrate. Thus, our data reveals transcriptional networks that govern MDSC developmental programs, and the impact of GCN2 stress signaling on the innate immune landscape in tumors, providing novel insight into potentially beneficial off target effects of dabrafenib.

## Introduction

Myeloid-derived suppressor cells (MDSCs) are immature monocytic and polymorphonuclear (PMN)-lineage cells that are expanded and activated in response to malignancy (Bronte et al., 2016). In cancer, MDSCs are defined by their potent ability to suppress T cell activation and function, contributing to immune suppression and cancer progression. In the most widely accepted model of MDSC development, PMN and monocytic MDSCs (m-MDSC) are derived from haematopoietic progenitors that expand and acquire immune-suppressive function in response to tumor-driven emergency granulopoiesis. Supporting this developmental model, *in vitro* analysis has shown that addition of inflammatory cytokines promoted development of MDSCs with potent ability to suppress T cell proliferation and effector function (Marigo et al., 2010). However, increased insight regarding the impact of tissue microenvironmental cues, stress signaling, the inflammatory landscape (local and systemic), and epigenetics on MDSC development/function has highlighted the complexity of MDSC biology, which is clearly still poorly understood (Hegde et al., 2021).

One important factor that influences MDSC physiology is intracellular and microenvironmental stress. For example, unfolded protein response (UPR) stress signalling transmitted by protein kinase R-like endoplasmic reticulum kinase (PERK) has a profound impact on MDSC suppressive function (Mohamed et al., 2020). Similarly, our group has reported that General Control Nonderepressible 2 (GCN2) is a key driver of MDSC function in the tumor microenvironment. Genetic deletion of *Eif2ak4* (the gene encoding GCN2) in myeloid cells altered MDSC metabolism and suppressive function enhancing anti-tumor CD8^+^ T cell immunity *in vivo* (Halaby et al., 2019). We also have found that GCN2 activation modulated macrophage and dendritic cell function in inflammatory disease altering cytokine production in endotoxemia (Liu et al., 2014a) and driving acquisition of a suppressive phenotype in autoimmunity (Ravishankar et al., 2015). The above findings suggested that relative GCN2 activity is an important modulating feature of myeloid biology regulating inflammatory and tolerogenic potential.

GCN2 is an evolutionarily ancient Ser/Thr kinase found in all eukaryotes that is an integral part of the integrated stress response (ISR), a cellular response system activated by diverse nutritional and environmental stress signals (Harding et al., 2000). GCN2’s kinase activity is induced by ribosomal stalling on mRNA transcripts resulting from paucity of aminoacyl-charged tRNA from amino acid starvation or active translation (Harding et al., 2019; Inglis et al., 2019). Once activated, GCN2 phosphorylates eukaryotic initiation factor (eIF)2α, significantly slowing GDP/GTP exchange in the translational complex abrogating cap-dependent translation (McGaha et al., 2012). Slowed ribosomal assembly resulting from reduced GTP priming by eIF2α also induces a transcriptional stress response increasing translation of activating transcription factor 4 (ATF4) and ATF5 which drives expression of amino acid transporters and modulates metabolism, autophagy and proliferation altering cellular phenotype and promoting survival (McGaha et al., 2012).

It has become increasingly clear that chemotherapeutics, radiation, and targeted therapies have the potential to control cancer growth by off-target effects impacting the cancer cells themselves and/or the induction of anti-cancer immunity (Wargo et al., 2015). In this vein, a growing body of literature suggest that target promiscuity by small molecule kinase inhibitors impacting GCN2 activity contribute to their therapeutic efficacy. For example, neratinib, an epidermal growth factor receptor (EGFR) inhibitor, activates GCN2 in glioblastoma cells contributing to its anti-tumor activity (Tang et al., 2022). Similarly, a kinome screen identified the EGFR inhibitor erlotinib and the receptor tyrosine kinase inhibitor sunitinib as agonists of GCN2 (Tang et al., 2022). Dabrafenib is a small molecule kinase inhibitor used to treat BRAF-mutant (mut) cancers, including melanoma and non-small cell lung cancer (Planchard et al., 2022; Robert et al., 2019). Mass spectrometry-based chemical proteomics analysis of dabrafenib target profiles in melanoma cells identified a putative interaction with GCN2, a predicted property not shared by another BRAF inhibitor, vemurafenib (Phadke et al., 2018). This interaction was confirmed in a chemical screen that identified dabrafenib as a potent GCN2 agonist (Li et al., 2018).

Since we have previously shown that GCN2 is essential for MDSC phenotype impacting metabolism and immune function, in this study we examined if dabrafenib-mediated modulation of GCN2 activity would impact MDSC development and function. Here, we show that direct activation of GCN2 by dabrafenib induced changes in MDSC differentiation blocking expansion and differentiation into PMN lineage cells. This altered development reshapes function by driving cells to acquire a phenotype with reduced suppressive capability.

## Results

### Dabrafenib impairs MDSC proliferation and differentiation

To study the impact of dabrafenib-GCN2 interaction on MDSC development, we utilized an *in vitro* model where bone marrow is cultured in the presence of GM-CSF and IL-6 for 4 days to generate immune suppressive MDSCs (Marigo et al., 2010) (Halaby et al., 2019). First, we assessed the dabrafenib effect on MSDC suppressive function. Enriched splenic CD11c^+^ dendritic cells (DCs) were pulsed with ovalbumin peptide OVA _a_nd co-cultured with transgenic T cells reactive against OVA_257-264 [_i.e. OTI CD8^+^ T cells (Hogquist et al., 1994)] in the presence or absence of MDSCs generated in a gradient of dabrafenib. Antigen pulsed DCs drove robust T cell proliferation (**Fig 1A** and **1B**), whereas addition of MDSCs reduced overall T cell proliferation by 60%, increasing the fraction of undivided cells from 5% to 40% (**Fig 1A** and **1B**). Strikingly, MDSCs generated in a gradient of dabrafenib showed a dose-dependent decrease in the ability to suppress T cell proliferation, indicating dabrafenib reduced MDSC suppressive activity. Increased enzymatic arginine metabolism is a hallmark feature of MDSC biology and suppressive function (Gabrilovich and Nagaraj, 2009). Dabrafenib treatment significantly decreased expression of the arginine metabolizing enzymes such as arginase 1 (*Arg1*) and a complete abrogation of inducible nitric oxide synthase (*Nos2*) (**Fig 1C**). This data shows that dabrafenib had a profound impact on the MDSC immune-suppressive phenotype.

**Figure 1.**
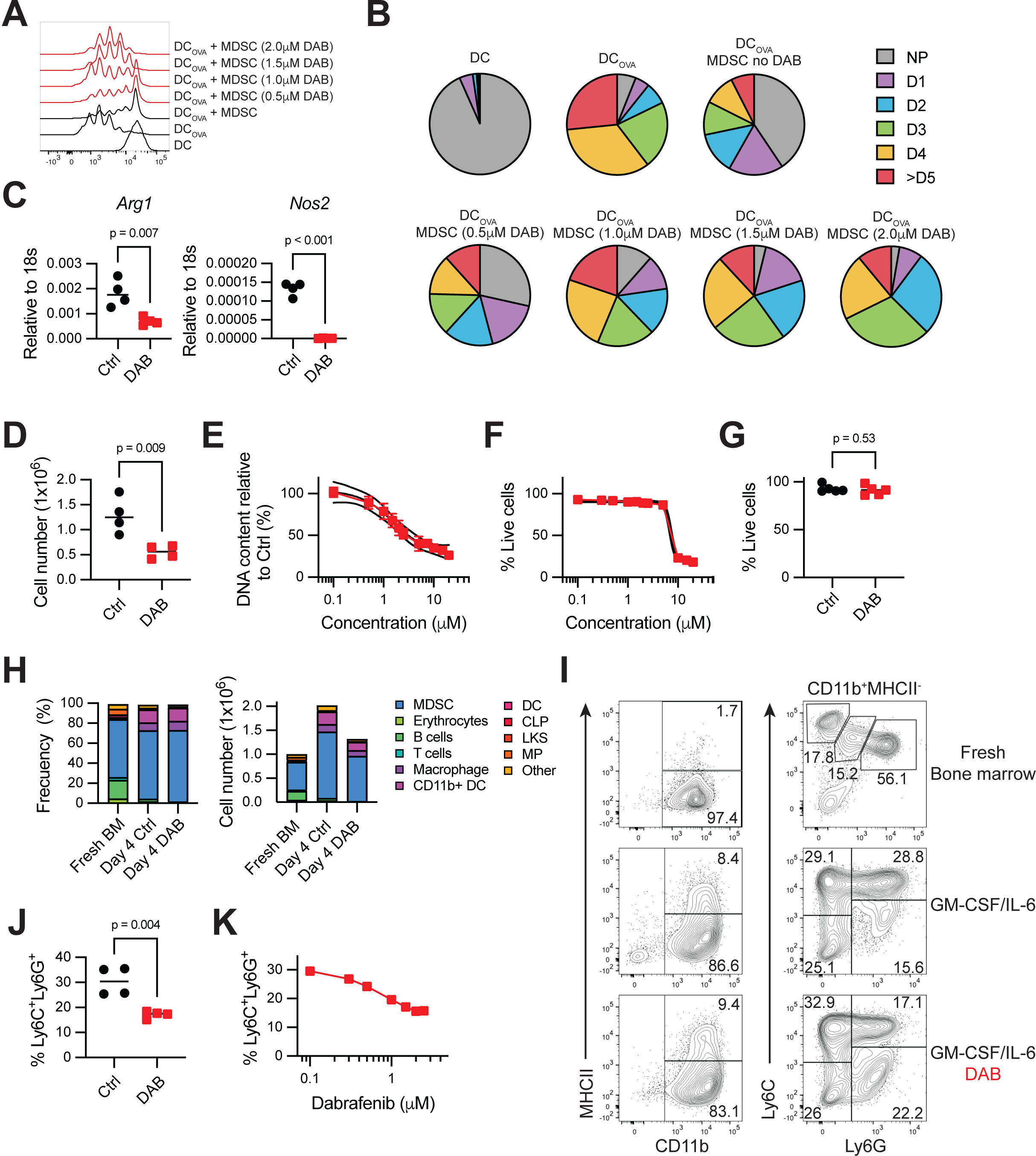
Dabrafenib (DAB) impairs MDSC proliferation and differentiation. **(A)** Representative flow cytometry histograms showing antigen-specific proliferation of CD8^+^ T cells from OTI mice modulated by bone marrow-derived MDSC. T cells were co-cultured with OVA_257-264 p_eptide-pulsed dendritic cells (DC) and MDSC at ratio DC: T cell: MDSC 1:10:30, where 1 = 10,000 cells, in 96-well plates for 60h. Experiment repeated 3 times with similar results. DAB=dabrafenib. **(B)** Frequency of OTI CD8+ T undergoing one (D1), two (D2), three (D3), four (D4) or more than five (D5) cycles of division from plot (A). NP=non-proliferative cells. Data are expressed as mean value. Experiment repeated 3 times with similar results. **(C)** Mean *Arg1* and *Nos2* expression relative to 18s RNA, assessed by quantitative real-time PCR (qPCR) (*n*=4). MDSCs were generated in presence ± 1.5μM DAB for 4 days. p-value was determined by Two-tailed unpaired t-Student’s test. Significance considered p<0.05. Experiment repeated 3 times with similar results. **(D)** Number of MDSCs generated *in vitro* in the presence of ±1.5μM DAB after 4 days in culture (n=4). p-value was determined by Two-tailed unpaired t-Student’s test. Significance considered p<0.05. Experiment repeated 3 times with similar results. **(E)** Dose-response for inhibition of proliferation after 4 days in culture. DNA fluorescence was measured for cultures with the indicated concentration of dabrafenib. IC50 value was calculated at 1.59μM. Lines represent the mean ± SEM of 4 independent experiments. **(F)** Dose-response for viability after 4 days in culture in presence of 1.5μM DAB. Each data point indicates the percentage of total live cells assessed by flow cytometry, normalized to control samples. Lines represent the mean ± SEM of 4 independent experiments. **(G)** Percentage of live cells in MDSC *in vitro* cultures ± 1.5μM DAB assessed by flow cytometry. Error bars indicate ± SD (*n*=4). p-value was determined by Two-tailed unpaired t-Student’s test. Significance considered p<0.05. Experiment repeated 4 times with similar results. **(H)** Mean frequency and total cell number of different subsets of immune cells in fresh bone marrow (BM) and MDSC. 1x10^6^ BM cells were seeded at the start of the experiment and differentiated for 4 days ± 1.5μM DAB (*n*=4). Color coding represents each subtype. Experiment repeated 2 times with similar results. **(I)** Representative flow cytometry contour plots of MDSC surface phenotype when expanded for 4 days with ± 1.5μM DAB compared to the ones found in fresh bone marrow. Gates show mean frequency (*n*=4). Experiment repeated 2 times with similar results. **(J and K)** Mean frequency of the PMN population CD11b^+^MHCII^neg^Ly6C^+^Ly6G^+^ in MDSC ± 1.5μM DAB (J) or increasing doses of DAB (K) (*n*=4). Experiment repeated 3 times with similar results.In panel **J**, p-value was determined by Two-tailed unpaired t-Student’s test. Significance considered p<0.05.

We then tested if the reduced MDSC suppressive function in the presence of dabrafenib was the result of toxicity. We observed that the higher concentrations of dabrafenib reduced overall numbers of MDSCs (**Fig 1D**). DNA content measurements indicated the IC50 for this effect was 1.59 μM (SEM±0.1) (**Fig 1E**). To determine if dabrafenib reduced cell numbers via toxicity, we quantified viability by flow cytometry. Cellular viability remained stable at ∼89% (SEM±2.61) for dabrafenib concentrations below 10 μM (**Fig 1F**). Importantly, when using a dabrafenib concentration of 1.5 ΙM (that reduced DNA content by 50%, **Fig 1E**), viability was not compromised, suggesting the effect was not due to cell death (**Fig 1G**).

We then asked if dabrafenib impacted MDSC expansion or cellular composition (**sFig 1**). In freshly isolated bone marrow, most cells expressed the general myeloid marker CD11b^+^ and a Ly6C^+^Ly6G^neg^ or Ly6C^+^Ly6G^+^ surface phenotype (57.8%, SEM± 1.5). Ly6C^+^Ly6G^+^ cells represented the majority (56.1%, SD±2.9) (**Fig 1H** and **1I**). In IL-6+GM-CSF expanded MDSC, CD11b^+^ cells were 94.3% (SD±1.8) of the total population after 4 days of culture (**Fig 1H** and **1I**). 1.5 1M dabrafenib did not have a significant effect on overall percentages of mature (CD11b^+^MHCII^+^) or immature (CD11b^+^MHCII^neg^) cells (**Fig 1H** and **1I**). However, PMN-MDSCs (Ly6C^+^Ly6G^+^) were significantly reduced from 28.8% (SD±5.1) to 17.1% (SD±4.5, p=0.004) (**Fig 1I** and **Fig 1J**). Dabrafenib-mediated reduction in the percentage of Ly6C^+^Ly6G^+^ cells was dose dependent (**Fig 1K**), paralleling the decrease in DNA content observed (**Fig 1E**) suggesting dabrafenib may impact PMN-MDSCs in a selective fashion.

### PMN MDSCs develop from monocytes and immature progenitors

Based on the above findings we predicted that dabrafenib may impact expansion of PMN-MDSC precursors. To test this, we labeled total bone marrow with a proliferation tracer dye prior to the addition of IL-6+GM-CSF, monitoring dye dilution over a 4-day culture period. Dabrafenib did not impact early proliferation of either the monocytic (CD3^neg^CD19^neg^CD11b^+^MHCII^neg^Ly6C^+^) or PMN (CD3^neg^CD19^neg^CD11b^+^MHCII^neg^Ly6C^+^ Ly6G^+^) populations; however, proliferation was significantly inhibited at day 4 for both monocytic and PMN populations with the PMNs more severely impacted by dabrafenib (**Fig 2A**). The monocytic population was actively proliferative with >90% of the cells exhibiting low amounts of tracer dye by day 4 (**Fig 2A**); in contrast PMN cells showed 2 distinct CSFE peaks on day 4. While the majority (61.6%, SD±6.1) showed a proliferative phenotype with low levels of tracer dye, 38.4% (SD±6.1) appeared quiescent without further cell division compared to day 1 (**Fig 2A**). Dabrafenib treatment significantly expanded this quiescent PMN population at day 4 to 67.9% (SD±6.2) of the population while a minority of the PMN cells now appeared proliferative (32.1%, SD±6.2) (**Fig 2A**). The subset of PMN cells that were quiescent on day 4 exhibited a surface phenotype distinct from proliferative PMNs, with decreased Ly6C expression (**Fig 2B**), likely forming the Ly6G^+^Ly6C^neg^ population observed when developing MDSCs were exposed to dabrafenib (**Fig 1I**). However, analysis of absolute cell numbers showed that GM-CSF+IL-6 preferentially expanded the monocytic population (Ly6C^+^) 13.0-fold ±4.7 compared to fresh bone marrow (**Fig 2C**). Despite being more numerous at culture initiation, Ly6G^+^Ly6C^+^ cells showed significantly less proliferative potential, expanding 2.2-fold ±1 relative to baseline numbers (**Fig 2C**). Dabrafenib exposure significantly reduced Ly6C^+^ cell expansion, but some proliferative potential remained as we observed a 2.0-fold expansion in Ly6C^+^ cells compared to starting numbers **(Fig 2C**). In contrast, Ly6G^+^ cells were severely impacted by dabrafenib and contracted by 50% at day 4 (**Fig 2C**). These data suggest the reduction in cellularity by dabrafenib exposure was caused, at least partially, by arrested Ly6C^+^ cell expansion, and indicated that PMN-MDSCs (CD11b^+^MHCII^neg^Ly6C^+^Ly6G^+^) might have originated from monocytic lineage cells.

**Figure 2.**
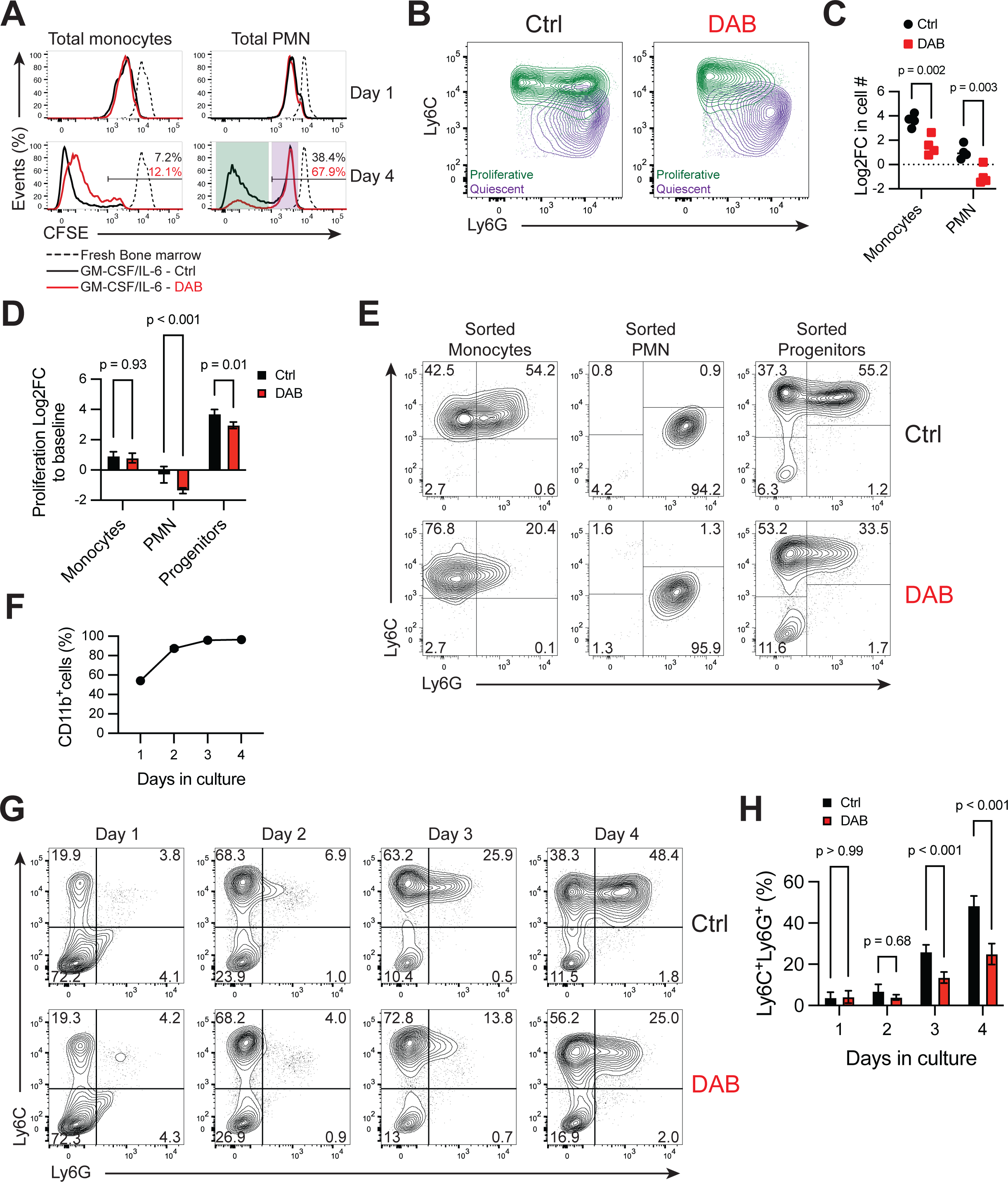
PMN-MDSCs develop from monocytes and immature progenitors. **(A)** Representative flow cytometry histogram of CFSE dilution in total monocytes (CD11b^+^MHCII^neg^Ly6C^+^) and total PMN (CD11b^+^MHCII^neg^Ly6G^+^) on day 1 and 4 in culture ± 1.5μM DAB. Day 4 panels show percent (%) of CFSE^high^ cells. The purple shaded area shows PMN cells that remain quiescent and do not proliferate beyond the initial cycle on day 1. The green shaded area shows PMN cells that are highly proliferative. **(B)** Representative flow cytometry contour plot of Ly6C and Ly6G expression in total PMNs gated as CD11b^+^MHCII^neg^Ly6G^+^ on day 4 after differentiation ± 1.5μM DAB. Quiescent (purple) and proliferative (green) PMN cells described in A are overlaid for each condition. **(C)** Log2 fold-change (Log2FC) of cell counts of total monocytes (CD11b^+^MHCII^neg^Ly6C^+^) and total PMNs (CD11b^+^MHCII^neg^Ly6G^+^) on day 4 normalized to the seeding counts of each lineage group at basal time point. MDSC were generated in presence of ± 1.5μM DAB. p-values were determined by Two-way ANOVA with Šidák correction post-test. Significance considered p<0.05. **(D)** Log2FC in proliferation measured by incorporation of DNA intercalating fluorescent dye on day 4, normalized to the fluorescence signal of cells at baseline. Sorted monocytes (CD3^neg^CD19^neg^CD11b^+^MHCII^neg^Ly6C^+^LyG6^neg^), PMN (CD3^neg^CD19^neg^CD11b^+^MHCII^neg^Ly6C^med^LyG6^+^) or progenitors (CD5^neg^CD11b^neg^CD19^neg^ B220^neg^Gr-1^neg^TER119^neg^) were differentiated to MDSC ± 1.5μM DAB. Cells were sorted from fresh bone marrow and plated at the same cell density (2x10^4^ cells/well). Bars show mean ± SD (*n*=4). p-values were determined by Two-way ANOVA with Šidák correction post-test. Significance considered p<0.05. **(E)** Representative flow cytometry contour plot of Ly6C and Ly6G expression in CD11b^+^MHCII^neg^ cells differentiated for 4 days. Initial cell source was either sorted bone marrow monocytes (CD3^neg^CD19^neg^CD11b^+^MHCII^neg^Ly6C^+^LyG6^neg^), bone marrow PMNs (CD3^neg^CD19^neg^CD11b^+^MHCII^neg^Ly6C^med^LyG6^+^) or bone marrow progenitors (CD5^neg^CD11b^neg^CD19^neg^B220^neg^Gr-1^neg^TER119^neg^). Gates show mean frequency. **(F)** Frequency of total CD11b^+^ cells during sorted progenitor differentiation, assessed by flow cytometry for each day of cell differentiation. Line represents mean ± SD (*n*=4). **(G)** Representative contour plot of Ly6C and Ly6G expression in CD11b^+^MHCII^neg^-cells each day during sorted progenitor differentiation. Gates show mean frequency. Experiment repeated 3 times with similar results. **(H)** Frequency of the granulocytic population CD11b^+^MHCII^neg^Ly6C^+^Ly6G^+^ during progenitor differentiation, determined by flow cytometry for each day of cell differentiation. Bars indicate mean ± SD (*n*=4). p-values were determined by Two-way ANOVA with Šidák correction post-test. Significance considered p<0.05. For all panels, experiments were repeated 3 times with similar results

MDSCs exhibit plasticity in expansion and differentiation potential, promoting MDSC contribution to multiple mature myeloid populations in tissue microenvironments. Thus, to elucidate the cellular dynamics contributing to population development, we examined the effect of dabrafenib on expansion and differentiation of FACS-enriched myeloid precursors or more mature PMN and monocytic lineage cells. Sorted progenitors (Lineage negative: CD5, CD11b, CD19, CD45R/B220, Ly6G/C(Gr-1), TER119, 7-4), monocytic (CD3^neg^CD19^neg^Ter119^neg,^ NK1.1^neg^CD11b^+^MHCII^neg^Ly6C^+^) or PMN (CD3^neg^CD19^neg^Ter119^neg,^ NK1.1^neg^CD11b^+^MHCII^neg^Ly6C^+^ Ly6G^+^) cells from bone marrow were seeded at the same cell density and cultured for 4 days with GM-CSF+IL-6 ±dabrafenib. Progenitors exhibited the greatest proliferative potential, expanding 6.9-fold more than enriched monocytic cells and 16.2-fold more than PMN cultures (**Fig 2D**). Monocytic cells nearly doubled (Log2FC = 0.9, SD±0.3), in stark contrast to isolated PMN populations that failed to expand in culture (**Fig 2D)**, indicating that the PMN accumulation observed in bulk cultures developed from progenitors and/or monocytic cells. Contrary to observations in bulk cultures, dabrafenib did not affect monocyte proliferation; however, dabrafenib accentuated PMN population contraction (**Fig 2C** and **2D**). Dabrafenib significantly reduced progenitor cell expansion (p<0.01), suggesting that differential progenitor sensitivity to dabrafenib may be driving the observed effects on overall MDSC expansion (**Fig 2D**). In progenitor cultures, exposure to IL-6+GM-CSF promoted development of monocytic Ly6C^+^Ly6G^neg^ and PMN Ly6C^neg^Ly6G^+^ MDSC populations (**Fig 2E**). Similarly, monocytic cultures showed emergence of a Ly6C^+^Ly6G^+^ MDSC population after culture with GM-CSF+IL-6 for 4 days (**Fig 2E**). Importantly, dabrafenib significantly reduced the Ly6C^+^Ly6G^+^ population for both monocytic and progenitor cultures (**Fig 2E**). This contrasted Ly6C^neg^Ly6G^+^ PMN cultures which lacked the ability to differentiate into Ly6C^+^ MDSCs in the presence or absence of dabrafenib (**Fig 2E**). Thus, the data suggests that dabrafenib limits differentiation plasticity of myeloid progenitors and newly generated mMDSCs.

To better understand the impact of dabrafenib on MDSC development from progenitors, we monitored the kinetics of MDSC emergence from precursor populations. Total CD11b^+^ cells comprised 54% (SD±3) of all cells on day 2, and significantly increased up to 95.9% (SD±0.6) by day 3 (p<0.001) (**Fig 2F**) an effect not impacted by dabrafenib. Monocytic Ly6C^+^Ly6G^neg^ MDSCs emerged as early as day 1 of culture, expanding to 68% (SD±4.3) of the total cultures by day 2. (**Fig 2G**). In contrast, PMN-MDSCs did not significantly expand until culture day 3 where they were 25.9% (SD±3.6) of all MDSCs, further expanding to 48.4% (SD±4.7) on day 4 (**Fig 2G**). Importantly the Ly6G^+^ population emerged from the Ly6C^+^ myeloid population (**Fig 2G**), consistent with the notion that PMN-MDSCs develop from the m-MDSCs. Dabrafenib did not impact m-MDSC expansion; however, PMN-MDSC expansion was significantly reduced (p<0.001) on day 3 and the PMN-MDSCs that developed exhibited lower overall Ly6G expression (**Fig 2G** and **2H**). Thus, the data suggests inflammatory cytokines drive sequential evolution of MDSCs into Ly6C^+^ populations that lead to later emergence of PMN-MDSCs. This transition from monocytic Ly6C^+^Ly6g^neg^ to PMN Ly6C^+^Ly6G^+^ MDSCs is attenuated by dabrafenib.

### Dabrafenib-induced GCN2 activation alters proliferation and differentiation of MDSCs

Dabrafenib targets BRAF-mut, however it also paradoxically hyperactivates the MAPK pathway in BRAF wild-type cells resulting in extracellular signal-regulated kinase (ERK)1 and ERK2 activation (Holderfield et al., 2014). Accordingly, we observed a dabrafenib dose-dependent increase in phospho-(p)ERK1/2 in MDSCs with a 2-fold induction over baseline phosphorylation at 1 1M (**Fig 3A** and **3B**). In control MDSCs, ERK1/2 activation was driven by a combination of IL-6 and GM-CSF signaling (**Fig 3C**), an effect that was significantly enhanced by dabrafenib. Decreased cellularity due to dabrafenib was observed only when GM-CSF was added to the cultures alone or in combination with other cytokines suggesting dabrafenib specifically impacts GM-CSF-induced cellular expansion (**Fig 3D**). Since dabrafenib enhanced ERK1/2 activation, we tested if this was responsible for altered differentiation of MDSCs *in vitro*. For this, we used the MEK inhibitor trametinib to block ERK1/2 phosphorylation in dabrafenib-exposed MDSCs. A trametinib concentration of 3 nM was sufficient to abrogate dabrafenib-induced p-ERK1/2 (**Fig 3E**). However, MDSC cultures treated with trametinib had reduced cell numbers comparable to dabrafenib-treated MDSCs (**Fig 3G**). Similarly, differentiation in presence of both dabrafenib and trametinib failed to restore dabrafenib-mediated PMN-MDSC phenotype (CD11b^+^MHCII^-^Ly6C^+^Ly6G^+^) observed in controls. (**Fig 3F**). This shows that increased MAPK activity was not responsible for dabrafenib-mediated effects on MDSCs.

**Figure 3.**
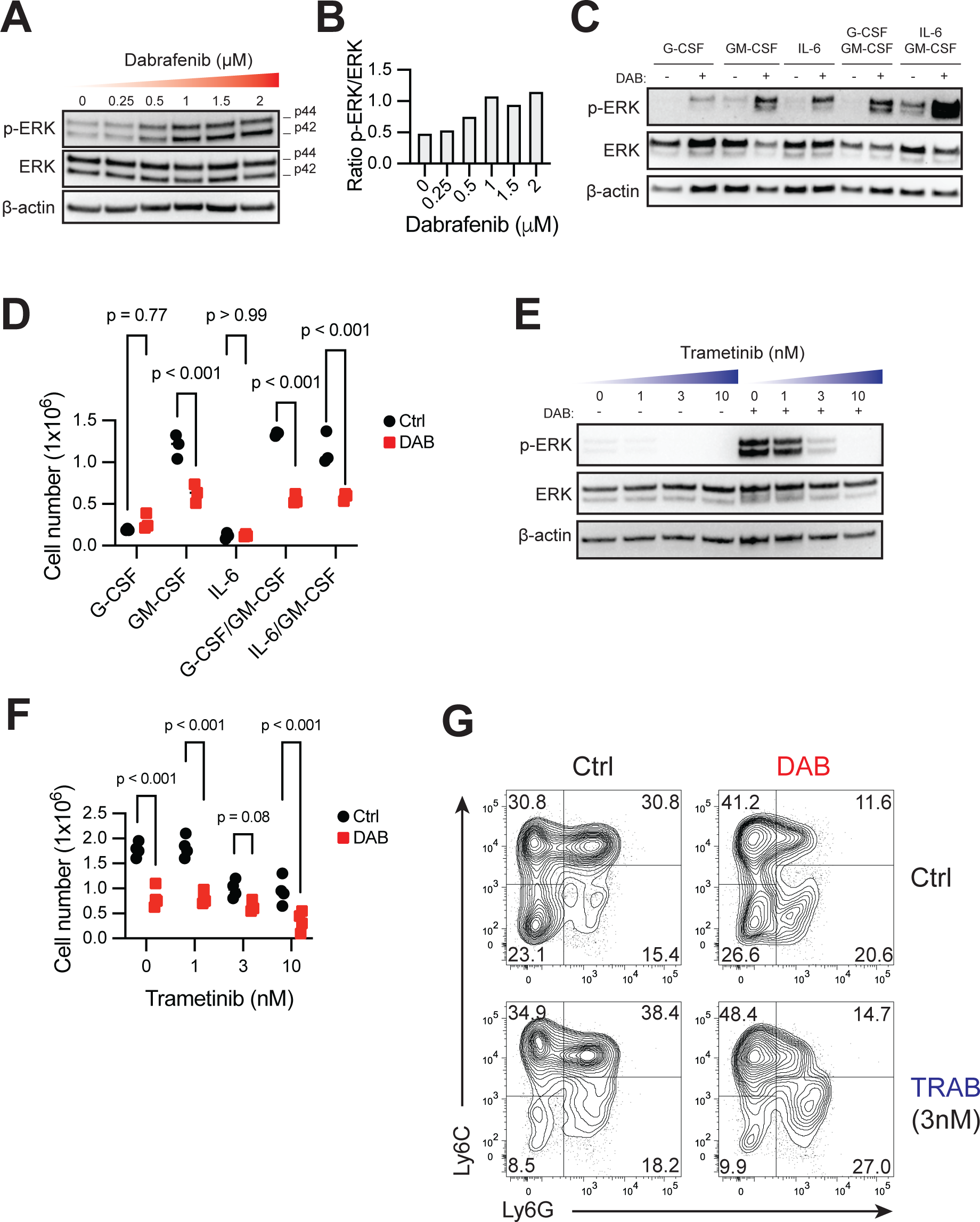
Activation of MAPK by Dabrafenib is not responsible for phenotypic changes during differentiation. **(A and B)** Western blot (**A**) and relative quantification (**B**) of ERK1/2 phosphorylation (p-ERK) versus total protein at increasing doses of DAB in bone marrow-derived MDSC cultures on day 4 of differentiation with IL-6+GM-CSF. **(C and D)** Western blot of p-ERK (**C**) and total cell counts (**D**) of bone marrow cells differentiated with indicated cytokines ± 1.5μM DAB for 4 days. For **D**, p-values were determined by Two-way ANOVA with Šidák correction post-test. Significance considered p<0.05. (**E and F**) Western blot of p-ERK (**E**) and total cell counts (**F**) of MDSC differentiated in presence of increasing doses of MEK inhibitor Trametinib (TRAB) ± 1.5μM DAB for 4 days. For **F**, p-values were determined by Two-way ANOVA with Šidák correction post-test. Significance considered p<0.05. **(G)** Representative contour plot of Ly6C and Ly6G expression in CD11b^+^MHCII^neg^ cells expanded in presence of 1.5μM DAB and/or 3nM TRAB. For all panels, experiments were repeated 3 times with similar results.

We then tested whether GCN2 activation was mechanistically required for dabrafenib modulation of MDSC differentiation. GCN2-deficient bone marrow was able to proliferate and develop into MDSCs in a manner comparable to GCN2 wild-type controls (**Fig 4A** to **4C**) suggesting GCN2 is not required for MDSC development. However, when GCN2 expression was lost, dabrafenib exposure no longer impacted expansion or differentiation of Ly6C^+^Ly6G^+^ MDSCs (**Fig 4A** to **4C**). Moreover, transcripts for the GCN2-responsive genes *Asns* and *Atf4* were significantly induced by dabrafenib, an effect that was completely abrogated in MDSC cultures lacking the gene coding GCN2, *Eif2ak4* (**Fig 4D**). This shows GCN2 is activated by dabrafenib and GCN2 function is required for dabrafenib’s MDSC inhibitory effects.

**Figure 4.**
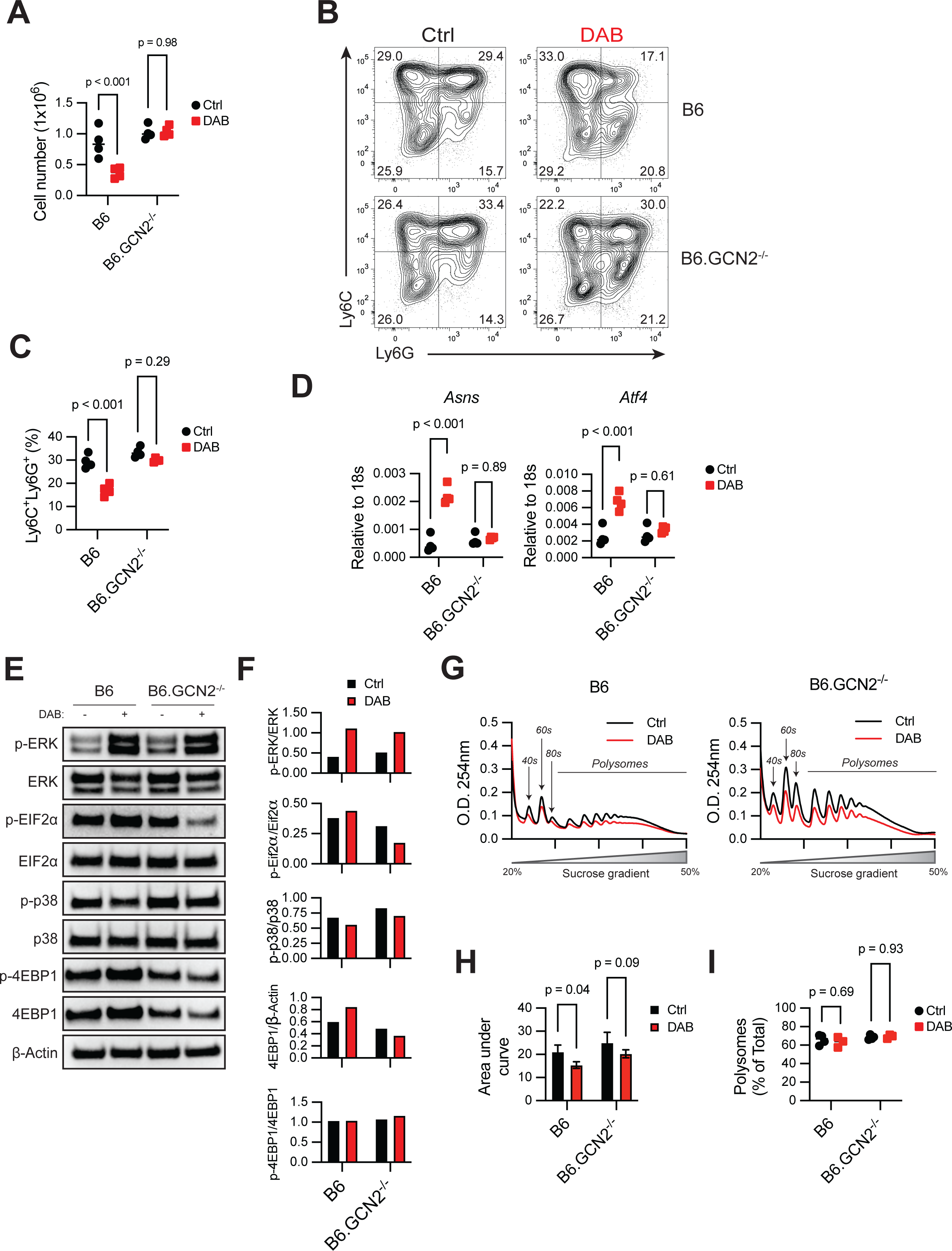
Dabrafenib-induced GCN2 activation alters proliferation and differentiation of MDSCs. **(A)** Total cell counts in MDSC *in vitro* cultures from wild-type (B6) and GCN2-deficient (B6.GCN2^-/-^) mice differentiated in presence of ± 1.5μM DAB. p-values were determined by Two-way ANOVA with Šidák correction post-test. Significance considered p<0.05. **(B)** Representative flow cytometry contour plot of Ly6C and Ly6G expression in wild-type and GCN2^-/-^ MDSCs differentiated in presence of ± 1.5μM DAB. **(C)** Frequency of CD11b^+^MHCII^neg^Ly6C^+^Ly6G^+^ MDSCs in wild-type and GCN2^-/-^ MDSCs differentiated in presence of ± 1.5μM DAB. p-values were determined by Two-way ANOVA with Šidák correction post-test. Significance considered p<0.05. **(D)** qPCR of *Asns* and *Atf4* in MDSCs lysates in wild-type and GCN2^-/-^ MDSCs differentiated in presence of ± 1.5μM DAB, relative to *18s* mRNA. p-value was determined by Two-tailed unpaired t-Student’s test. Significance considered p<0.05. **(E and F)** Western blot (**E)** and quantification (**F**) of ERK1/2, eIF2α, p38 and 4EBP1 phosphorylation relative to total protein in whole culture MDSC lysates. (**G to I**) Ribosomal profiling analyzed in wild-type and GCN2^-/-^ MDSC lysates. (**G**) Optical density (O.D.) at 254nm. The area designated as “polysomes” represents the fraction of RNA forming complexes of two or more ribosomes. (**H**) Area under the curve of the region marked as polysomes in **G**. Bars indicate mean ± SD (*n*=4). p-values were determined by Two-way ANOVA with Šidák correction post-test. Significance considered p<0.05. (**I**) Relative quantification (%) of the area under the curve in polysomes vs monosomes (40s, 60s and 80s region). Data are expressed as mean ± SD (*n*=4). p-values were determined by Two-way ANOVA with Šidák correction post-test. Significance considered p<0.05. For all panels, experiments were repeated 3 times with similar results.

We next examined how dabrafenib/GCN2 interaction impacted ISR responses in MDSCs. eIF2α, the main target of GCN2 kinase activity, showed constitutive phosphorylation in control MDSC cultures, likely as a result of proliferative and translational stress (**Fig 4E** and **4F**). Dabrafenib exposure increased phospho (p)-eIF2α suggestive of enhanced GCN2 activity (**Fig 4E** and **4F**). Loss of *Eif2ak4* did not impact basal p-eIF2α in MDSCs; however, addition of dabrafenib caused an unexpected 50% reduction in p-eIF2α in GCN2-deficient MDSCs compared to wild type controls (**Fig 4E** and **4F**). Moreover, in GCN2^-/-^ MDSCs there were reduced overall levels of 4EBP1, a negative regulator of translation (**Fig 4E**) (Qin et al., 2016). This suggested that in the absence of GCN2, dabrafenib reduced ISR signaling potentially enhancing translation. Thus, taken together the data suggests GCN2 is activated by dabrafenib, and GCN2 stress signaling is the principal mechanism that drives altered MDSC differentiation.

We have shown previously that GCN2 activation can alter ribosome association with mRNA in myeloid cells resulting in a shift from polysomes to single ribosome-bound mRNA, a hallmark of reduced protein translation activity (Ravishankar et al., 2015). Therefore, we examined if dabrafenib treatment impacted RNA/ribosome association in MDSCs in a GCN2-dependent manner. Sucrose gradient analysis of RNA/ribosome complexes showed that dabrafenib reduced overall association of ribosomes with mRNA (**Fig 4G** and **4H**). However, it did not change the relative frequency of polysomes versus 40s/60s subunits or 80s ribosomes suggesting dabrafenib did not negatively impact polysome formation in a specific manner (**Fig 4I**). Importantly, GCN2-deficient MDSCs exhibited a similar pattern of polysome association with mRNA (**Fig 4I**). However, the dabrafenib-induced reduction in overall ribosome association with mRNA transcripts was attenuated in GCN2-deficient MDSCs (**Fig 4G** and **4H**). Thus, the data suggest that dabrafenib impacts development of MDSCs by a GCN2-dependent mechanism, with a general reduction in ribosome/mRNA interaction.

### Dabrafenib induces broad transcriptional changes in MDSCs

GCN2-induced transcriptional responses are key drivers of its effect on myeloid cell phenotype (Halaby et al., 2019), therefore we next analyzed the transcriptomic responses in dabrafenib-treated MDSC subpopulations by single cell RNA sequencing. Most cells were identified as myeloid, expressing *Itgam* (the gene encoding CD11b), with varying expression of MHC molecules (e.g. *H2-Aa*) and lineage identity markers (e.g. *Ly6c2* and *Ly6g*) (**Fig 5A** and **5B**). Cluster identity analysis revealed an array of bone marrow enriched developmental cell types including immature monocyte and PMN precursors and more mature monocytic and PMN populations. Four clusters were transcriptionally similar to mature PMNs found in the peritoneal cavity (clusters 1, 5, 11 and 13) expressing *Cfs3r*, *Mmp9*, *Cd9*, and *Il1rn* (Jerome et al., 2022) (**sFig 2**). Additionally, five clusters (clusters 3, 4, 8, 9, and 10) were transcriptionally similar to developing PMNs found in the bone marrow, suggesting a more intermediate phenotype (Barman et al., 2022), while cluster 6 was transcriptionally similar to immature, Ly6G^lo^ neutrophils found in the peritoneal cavity (Jerome et al., 2022) characterized by expression of several immunomodulatory chemokines including *Ccl6*, *Cxcl2*, *Ccl3*, *Cxcl3*, and *Ccl4* (**sFig 2**). Three clusters (clusters 0, 2, and 12) identified as immature monocyte lineage cells characterized by expression of *Spp1*, *Lpl*, *Mgl2*, *Mmp12*, and *Ccr2* (**Fig 5B** and **sFig 2**). There were 2 clusters (cluster 7, 20) with a mature monocyte transcriptional profile including expression of genes related to antigen presentation (*H2-Aa, H2-Ab1*, *H2-Eb1* and *Cd74*) and migration and recruitment (*Ccr7*, *Ccl22*, and *Ccl5*) (**sFig 2**).

**Figure 5.**
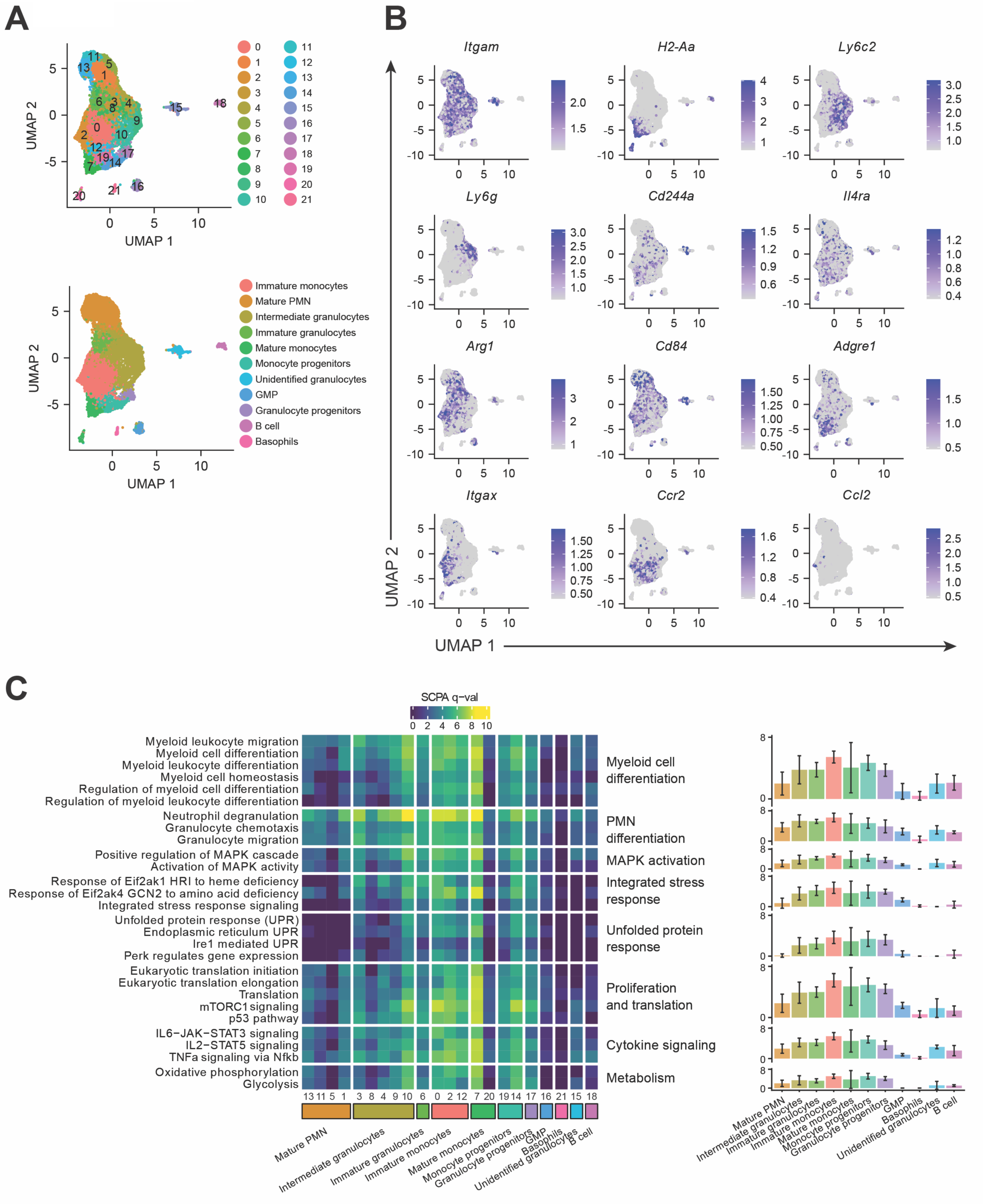
Dabrafenib induces broad transcriptional changes in MDSCs. **(A)** Upper-UMAP plot of the integrated scRNA sequencing data from MDSC differentiated ± 1.5μM DAB, showing the 22 clusters defined by Seurat analysis. Lower-UMAP plot of the integrated scRNA showing the 11 annotated lineage clusters defined by ProjecTILs (v3.0.0) and refined manually. **(B)** UMAP plot of the integrated scRNA sequencing data from MDSCs showing RNA expression of selected myeloid markers. Scale represents normalized RNA expression. **(C)** Heatmap of the SCPA multivariate distribution analysis of genes associated with the Hallmark, Reactome and GO Biological process pathways. Pathways are representative top biological processes involved in myeloid cell and PMN differentiation, MAPK activation, Integrated Stress Response (ISR), Unfolded Protein Response (UPR), proliferation and translation, cytokine signaling and respiration metabolism. Left panel shows q-val for individual pathways in each cluster. Right panel shows the average q-val ±SD for the pathways included in the biological process indicated, for each annotated group. Higher q-val represent pathways with greater differences upon stimulation with DAB versus control. q-val is defined by SCPA as the square-root of the log10(p-adjusted values) based on the Rosenbaum cross-match test.

To evaluate the overall impact of dabrafenib in each cluster, we performed multivariate distribution analysis with single cell pathway analysis (SCPA) for each cluster comparing dabrafenib to controls (Bibby et al., 2022). As expected, processes involved in myeloid and granulocyte differentiation were among the top regulated pathways in most clusters (adjusted p-val <0.01) as shown by a higher q-val statistic (**Fig 5C**). Importantly, higher q-val were observed in immature granulocyte and immature monocyte clusters and, to a lesser extent, precursor clusters supporting the prediction that dabrafenib modulates differentiation at stages where cells still possess plasticity. Moreover, MAPK pathway regulation and GCN2 response genes were also significantly influenced by dabrafenib while the UPR was mildly impacted **(Fig 5C**). Additionally, dabrafenib treatment modulated proliferation and translation as well as STAT3, STAT5, and NF-ΙB transcriptional modules in most clusters (**Fig 5C**).

### MDSC differentiation arrest is a result of a combination of cell cycle and myeloid lineage alteration

We identified the stage at which dabrafenib induced developmental arrest by evaluating the variation in cell frequency for each cluster. The relative frequency of cells in dabrafenib versus DMSO samples was calculated per each individual cluster using Pearson’s Chi-Square test where standardized residuals indicated deviation away from the null assumption of equal proportions and values greater than ±3 indicated a substantial differences (**Fig 6A**). Using this approach, we identified three intermediate clusters (cluster 4, 8, 9) and one mature PMN cluster (cluster 1) significantly reduced by dabrafenib (**Fig 6B**). Interestingly, the second most affected cluster was classified as immature monocytes (cluster 2) suggesting that monocyte subset differentiation is also compromised by dabrafenib (**Fig 6B**). Likewise, dabrafenib exposure was also associated with accumulation of mature PMN clusters 5, 11, 13 **(Fig 6B**). This indicates that dabrafenib primarily impacts the maturation state of PMN-MDSCs, reducing developmental intermediate populations in favor of more mature neutrophil populations.

**Figure 6.**
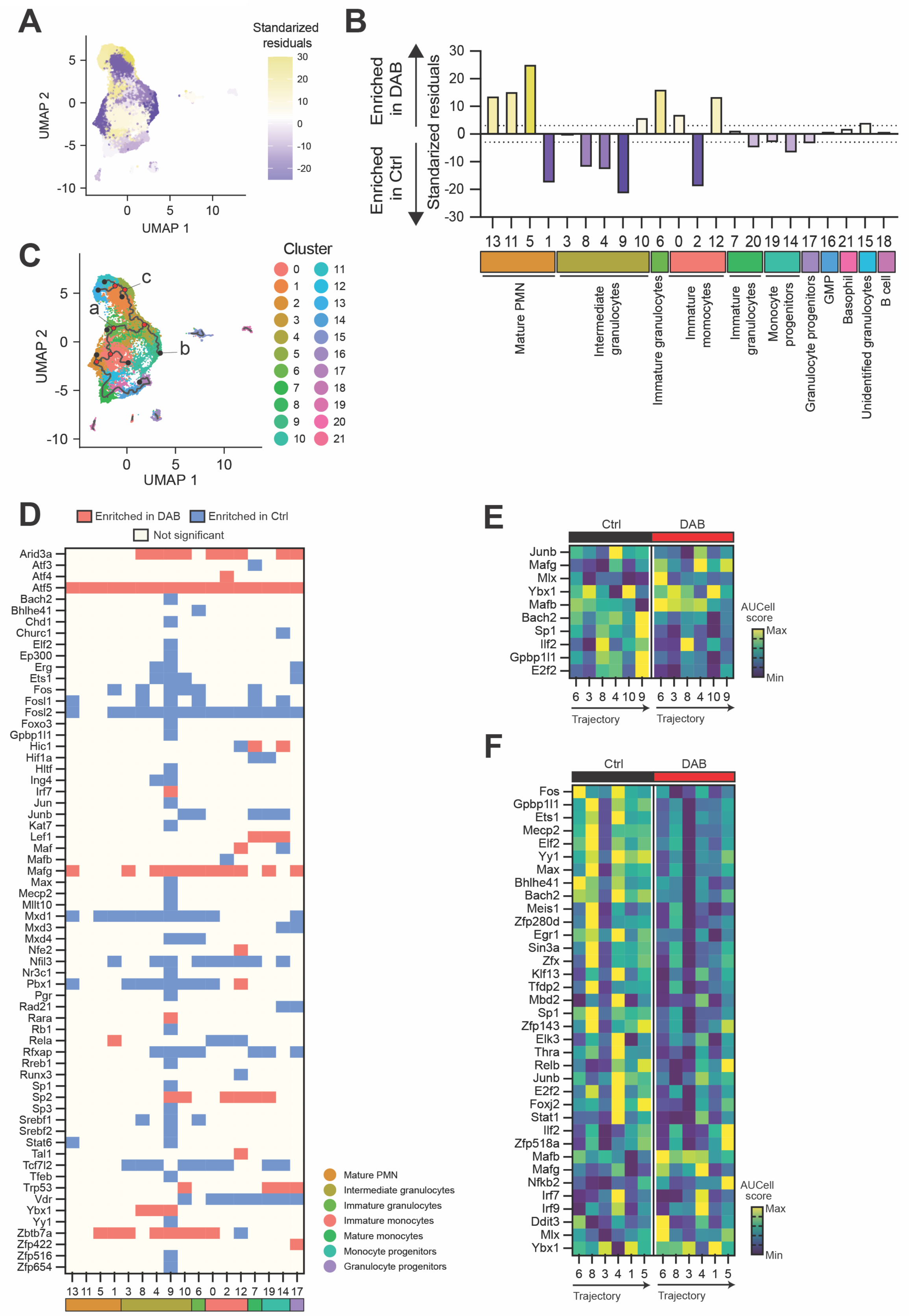
MDSC differentiation arrest is a result of a combination of cell cycle and myeloid lineage alteration. **(A and B)** Relative frequency of cells in each Seurat cluster from scRNA sequencing data of MDSCs ± 1.5μM DAB. Relative levels were calculated based on the standardized residuals from using Pearson’s Chi-Square overlaid on the UMAP plot of integrated samples (**A**) and grouped by functional annotation (**B**). Positive standardized residuals (yellow) for each cluster indicates increased frequency in the DAB treated sample whereas a negative number (purple) indicates increased frequency in the DMSO control sample. A dotted line indicates standardized residual values of ±3. **(C)** UMAP plot of the integrated scRNA sequencing data from MDSCs ± 1.5μM DAB, showing the trajectory branches calculated by Monocle3 and represented over the 22 clusters defined by Seurat. Most relevant nodes at the end of the branches (black dots) and at intersections (red dots) are shown. Nodes used as starting (a) or end (b and c) points for regulon analysis across the branch are labeled. **(D)** Heatmap of the regulons significantly regulated by DAB calculated on AUCell (q < 0.01, |Log2FC| >0.01) for each Seurat cluster. Color represents regulons significantly enriched in cells treated with DAB (red), those enriched in control samples (blue) and the ones that do not show significant changes (light yellow). **(E)** Heatmap of the regulons significantly regulated during differentiation in the direction from node a to node b in ***C***, which spans from cluster 6 (immature granulocytes) to cluster 9 (intermediate granulocytes) in DMSO control cell cultures. Scale represents z-score of the normalized average AUCell score per regulon. **(F)** Heatmap of the regulons significantly regulated during differentiation in the direction from node a to node c in ***C***, which spans from cluster 6 (immature granulocytes) to cluster 5 (mature PMN) in DMSO control cell cultures. Scale represents z-score of the normalized average AUCell score per regulon.

In contrast, PMN cluster 6 and monocytic cluster 12 were the only immature populations enriched by dabrafenib treatment (**Fig 6B**), suggesting they may represent transitional maturation stages where development is arrested influencing overall changes in cellular composition. To test this hypothesis, we performed trajectory analysis in the control DMSO sample which confirmed that cluster 6 is an early stage founding node for PMN lineage differentiation (**Fig 6C**). One branch followed the path towards more differentiated PMN stages while cluster 4 appeared to be a divergent node for an intermediate phenotype (cluster 9) or more mature stages (cluster 1, 5, 11, 13) (**Fig 6C**). The other branch connected to monocytic clusters and continued to progenitor clusters reinforcing the concept that such granulocytes derived from monocytic lineage cells. Taken together with the flow cytometry data, trajectory analysis suggests dabrafenib impacted MDSC differentiation at two stages: **1)** dabrafenib impedes the transition from monocyte-to-PMN in cluster 12, and; **2)** dabrafenib blocks early PMN differentiation from cluster 6 to more mature differentiation states.

To test this prediction, we performed downstream regulon analysis using the SCENIC package and calculated the AUCell enrichment scores for comparison between the treatment groups, focusing on the main group of clusters that showed interconnected trajectory branches. Like pathway analysis, we observed a generalized regulation of predicted transcription factor activity (q < 0.01, |Log2FC| >0.01). Dabrafenib induced enrichment of *Atf5*, *Mafg* and *Zbtb7a* regulons in the majority cell clusters (**Fig 6D, sFig 3A**). *Atf5* is directly regulated by GCN2 activation (Dang Do et al., 2009), and its enrichment in all clusters supports the prediction of broad, dabrafenib mediated, GCN2 induction. *Zbtb7a* regulates hematopoietic development and glycolysis, thus its activation is likely to have a significant impact on cellular metabolism and development (Liu et al., 2014b; Lunardi et al., 2013). Likewise, the Maf transcription factor family impacts myeloid cellular identity and maturation (Vanneste et al., 2023), and *MAFG* expression is most abundant in intermediate neutrophil precursors with significant downregulation in mature PMNs as they exit the bone marrow (Grassi et al., 2018). *Mxd1*, *Tcf7l2* and *Fosl2* regulons were enriched in control samples suggesting dabrafenib inhibited their activity. Importantly, decreased enrichment of the *Mxd1* regulon was found in PMN clusters preferentially in dabrafenib-treatment conditions (**Fig 6D, sFig 3A**). *Mdx1* codes for MAD1, a negative regulator of MYC and downstream ribosome biogenesis (Lafita-Navarro et al., 2016). Moreover, MAD1 binds DNA complexes during PMN differentiation (Ryan and Birnie, 1997; Walkley et al., 2004) and influences fitness and maturation of conventional dendritic cells (Anderson et al., 2020). Thus, a loss of the *Mdx1* regulon may be associated with reduced ribosomal fitness and differentiation potential, which would be in line with our observations.

Focusing on cluster 12 (immature myeloid), which represented a key point of cell accumulation, we identified four regulons that were exclusively regulated by dabrafenib in this monocytic cluster. Cluster 12 showed enriched *Nfe2* and *Tal1* regulons after dabrafenib treatment with decreasing *Runx3* regulon activity (**Fig 6D, sFig 3B**). TAL1 is a cell cycle activator in myeloid cells that inhibits expression of p21 and p16 (Dey et al., 2010) while RUNX3 is required for development and anti-inflammatory functions of mononuclear cells (Hantisteanu et al., 2020) providing a transcriptional basis for increased accumulation of cluster 12 (**Fig 6B**) and reduced overall T cell suppressive activity after dabrafenib exposure (**Fig 1A**). Additionally, we observed that PMN cluster 9 exhibited the highest number of downregulated transcriptional circuits with reduced enrichment of 36/44 regulons (**Fig 6D**), paralleling the significant decrease in cluster abundance after dabrafenib treatment (**Fig 6B**) suggesting these regulons are key drivers of PMN-MDSC development and function.

To better understand the influence of transcription factors programs on MDSC development ± dabrafenib, we calculated regulon enrichment in the trajectory branches identified. Since loss of PMN-MDSCs was the main effect of dabrafenib exposure, we focused on the importance of cluster 6 as the nascent population for developmentally intermediate PMN (cluster 9) and mature PMN-MDSCs (cluster 5). The trajectory analysis through cluster 6 to cluster 9 revealed *Bach2*, *Gpbp1l1* and *E2f2* are regulons that are progressively activated in the developmental trajectory to intermediate PMNs (**Fig 6E**). Those regulons were among the 22 regulons specifically modulated by dabrafenib in cluster 9 (**Fig 6D**) suggesting their activation was necessary to differentiate to an intermediate PMN phenotype. *Rb1* is a regulator of PMN differentiation in a recently characterized monocytic progenitor subtype of committed neutrophil precursors (Mastio et al., 2019). We identified *Rb1* regulon as significantly modulated by dabrafenib in cluster 9 (**Fig 6D**) but was not identified in the cluster 6 to cluster 9 trajectory branch (**sFig 3C**) suggesting that differential Rb was not driving PMN maturation in our samples.

We identified 36 regulons that were differentially regulated along the cluster 6-to-cluster 5 trajectory branch in control cultures (**Fig 6F**). Dynamic upregulation of their activity occurred during maturation progression and dabrafenib showed a broad inhibitory effect on regulon activity in general (**Fig 6F**). In contrast, the regulon controlled by MAFG was increased by dabrafenib in this differentiation branch. The *Mafg* regulon showed relatively low enrichment in intermediate PMN clusters in controls; however, in dabrafenib-treated cells, the *Mafg* regulon was significantly enriched (**Fig 6E** and **Fig 6F**). This suggests precise regulation of *Mafg* activity is essential for control of PMN-MDSC differentiation and dabrafenib-altered activity is likely a key driver of altered maturation/functional programs observed.

We have shown that GCN2 activity can alter MDSC metabolism, impacting MDSC function. Since several of the regulons identified can modulate metabolism and cellular energetics, we asked if dabrafenib alters MDSC metabolism by GCN2 activation. First, we examined expression of metabolism-associated genes in the MDSC single cell sequencing dataset, using a gene set we had previously identified as differentially expressed GCN2-deficient tumor macrophages and MDSC (Halaby et al., 2019). *Cox6a2* (encoding a subunit of cytochrome C oxidase), was strongly induced by dabrafenib in monocytic and progenitor clusters; however, there was no apparent induction in the more mature PMN clusters except for the key intermediate PMN cluster 9 (**Fig 7A).** Expression of genes involved in mitochondrial respiration (*Ppar©*), or glycolysis (*Aldh18a1*, *Slc2a1*, *Slc16a3*, *Ldha*) were also impacted by dabrafenib; however, the effect was less penetrant overall suggesting dabrafenib may enhance mitochondrial respiration (**Fig 7A**). Oxidative respiration is one of the major endogenous sources of reactive oxygen species (ROS) and can induce ribosome stalling and GCN2 activation. We measured ROS by staining MDSCs with CellROX for 1h before analysis. We observed that dabrafenib increased mitochondrial and nuclear ROS (**Fig 7B)**. This suggested that dabrafenib increased mitochondrial respiration in MDSCs. Supporting this, mRNA measurements in bulk dabrafenib-treated MDSC cultures also demonstrated strong induction of *Cox6a2*, an effect that was completely abrogated by loss of GCN2 function (**Fig 7C**). We then measured metabolic flux by seahorse, which showed that dabrafenib increased maximum oxidative respiration in the cultures 2-fold compared to controls but did not affect glycolytic flux (**Fig 7D** and **7E**). Moreover, in agreement with the mRNA analysis, loss of GCN2 function prevented dabrafenib-induced oxidative capacity increase **(Fig 7D** and **7E**). GCN2-deficient MDSCs showed a slight decrease in glycolysis overall, but glycolytic metabolic function was not impacted by dabrafenib (**Fig 7D** and **7E**). Thus, the data show that dabrafenib impacts metabolism, increasing oxidative respiration by activation of GCN2.

**Figure 7.**
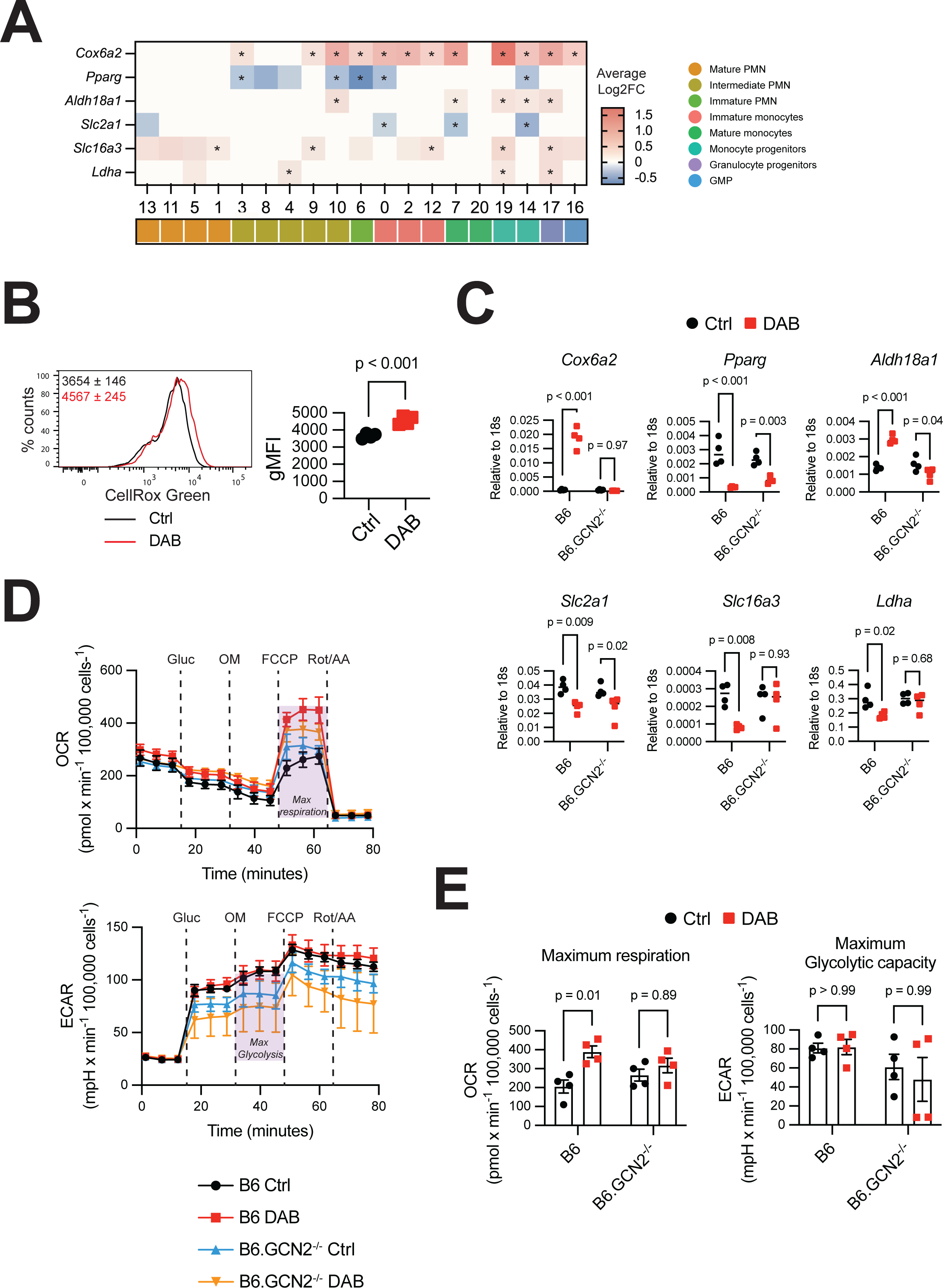
Dabrafenib increases oxidative metabolism in MDSCs. **(A)** Heatmap of the average Log2FC in gene expression for the genes indicated ± 1.5μM DAB clusters identified in MDSC scRNA sequencing data described in Fig 6. Positive values indicate genes with induced expression by DAB, while negative values show genes that are downregulated by DAB. For representation, Log2FC=0 indicates clusters that were excluded from the analysis with *FindAllMarkers* function (gene expressed in <10% cells and/or Log2FC ±0.25). Statistics show Wilcoxon test. * = adjusted p-value <0.05. **(B)** Representative flow cytometry histogram of CellROX Green (ROS in mitochondria and nucleus) in wild-type MDSC generated with1.5μM DAB (red line) or control (black line). Panels show geometric mean fluorescence intensity (gMFI) ±SD (*n*=4). p-value was determined by Two-tailed unpaired t-Student’s test. Significance considered p<0.05. Experiment was repeated 3 times with similar results. **(C)** qPCR of genes related to oxidative respiration (*Cox6a2*), fatty acid metabolism (*Pparg*) and glycolysis (*Aldh18a1, Slc2a1, Slc16a3, Ldha*) in total wild-type and GCN2^-/-^ MDSC lysates, normalized to *18s* mRNA expression(n=4). p-values were determined by Two-way ANOVA with Šidák correction post-test. Significance considered p<0.05. Experiment was repeated 3 times with similar results. **(D)** Quantification of oxidative respiration by oxygen rate consumption (OCR) and glycolysis by extracellular acidification rate (ECAR), by Seahorse assay. Lines represent the mean ±SEM of 4 independent experiments. Gluc = Glucose; OM = Oligomycin; FCCP = Carbonyl cyanide-p-trifluoromethoxyphenylhydrazone; Rot/AA = Rotenone/Antimycin **(E)** Maximum respiration calculated from OCR data and maximum glycolytic capacity calculated from ECAR data. Bars represent mean ± SEM of 4 independent experiments. p-values were determined by Two-way ANOVA with Šidák correction post-test. Significance considered p<0.05.

### Dabrafenib reduced MDSC accumulation in the Tumor Microenvironment

We then tested if dabrafenib would impact MDSC accumulation *in vivo* in the YUMM1.7 model of melanoma. YUMM1.7 possesses several of the hallmark driver mutations in melanoma including loss of *Pten* and *Cdkn2a* and the dabrafenib sensitive Braf^V600E^ mutation (Meeth et al., 2016). YUMM1.7 cells were implanted in the right hind leg and on day 10, mice were allocated into the treatment groups according to their tumor volume to reduce tumor size variability between groups. Mice were then treated for 7 days with 30 mg/kg dabrafenib and analyzed by flow cytometry for MDSC composition in the bone marrow, blood and tumor. In the bone marrow, dabrafenib treatment did not alter the number of live cells (**Fig 8A**); however, MDSC numbers were reduced by 50% compared to control mice (**Fig 8B**). In agreement with the *in vitro* data, 70% of the CD11b^+^MHCII^neg^ cells in the bone marrow were Ly6C^int^Ly6G^+^ PMN lineage cells (**Fig 8C** and **8D**). Although there was no reduction in m-MDSC versus PMN-MDSC populations after dabrafenib treatment (ratio PMN/m-MDSC 5.5±1.7 in Ctrl and 6.0±0.6 in DAB); **Fig 8D**), the dominance of PMN lineages in the MDSC gate (66.4%±5.1 and 69.6%±2.5 in Ctrl and DAB, respectively) suggests these cells are impacted by dabrafenib in agreement with our *in vitro* observations.

**Figure 8.**
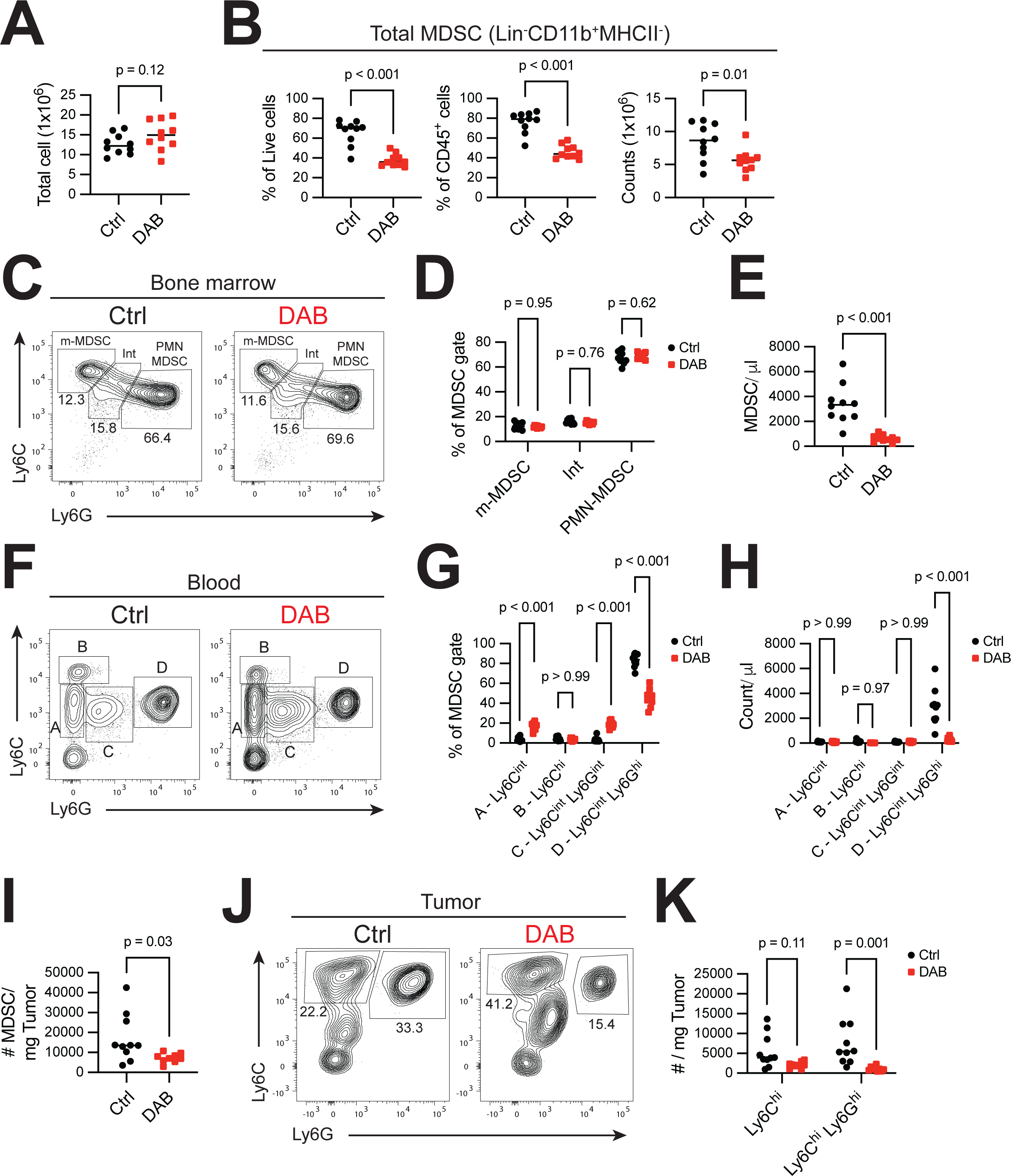
Dabrafenib reduces MDSC accumulation in the tumor microenvironment. Mice were implanted with 3x10^5^ YUMM1.7 cells subcutaneously in the right flank. On Day 10 after tumor implantation mice were either treated with 30mg/kg of DAB or the equivalent volume of DMSO for 7 consecutive days (*n*=10 per group). Mice were sacrificed on day 20 after tumor implantation. **(A)** Number of total live cells in bone marrow, isolated from one tibia and femur per mouse after red cells lysis. p-value was determined by Two-tailed unpaired t-Student’s test. Significance considered p<0.05. **(B)** Frequency of MDSCs (CD45^+^CD3^neg^CD19^neg^CD11b^+^MHCII^neg^) from total live cells (left panel), from immune cells (middle panel), and absolute counts of MDSCs (right panel). p-value was determined by Two-tailed unpaired t-Student’s test. Significance considered p<0.05. **(C)** Representative contour plot of Ly6C and Ly6G expression in total MDSCs (CD45^+^CD3^neg^CD19^neg^CD11b^+^MHCII^neg^) in bone marrow. Gates show median frequency of m-MDSCs (Ly6C^+^Ly6G^neg^), PMN-MDSCs (Ly6C^+^Ly6G^+^) and cells with intermediate phenotype (Int, Ly6C^int^Ly6G^int^). **(D)** Frequency of MDSCs subtypes described in ***C***. p-values were determined by Two-way ANOVA with Šidák correction post-test. Significance considered p<0.05. **(E)** Number of circulating MDSCs (CD45^+^CD3^neg^CD19^neg^CD11b^+^MHCII^neg^) per microliter of blood. **(F)** Representative contour plot of Ly6C and Ly6G expression in total MDSCs (CD45^+^CD3^neg^CD19^neg^CD11b^+^MHCII^neg^) in blood. Gates show simplified nomenclature for all four subpopulations a) Ly6C^int^ b) Ly6C^hi^ c) Ly6C^int^Ly6G^int^ and d) Ly6C^int^Ly6G^hi^. **(G)** Frequency of MDSC subtypes described in ***F***. p-values were determined by Two-way ANOVA with Šidák correction post-test. Significance considered p<0.05. **(H)** Absolute counts of MDSC subtypes described in ***F*** and ***G*** per microliter of blood. p-values were determined by Two-way ANOVA with Šidák correction post-test. Significance considered p<0.05. **(I)** Number of intratumoral MDSCs (CD45^+^CD11b^+^MHCII^neg^) per milligram of tumor. p-value was determined by Two-tailed unpaired t-Student’s test. Significance considered p<0.05. **(J)** Representative contour plot of Ly6C and Ly6G expression in total intratumoral MDSCs (CD45^+^CD3^neg^CD19^neg^NK1.1^neg^CD11c^neg^CD11b^+^MHCII^neg^). Gates show median frequency of monocytic MDSCs (Ly6C^hi^) and PMN-MDSCs (Ly6C^hi^Ly6G^hi^). **(K)** Absolute counts of MDSC subtypes described in ***J*** per milligram of tumor. p-values were determined by Two-way ANOVA with Šidák correction post-test. Significance considered p<0.05. Panels are representative of 3 independent experiments.

MDSCs (CD45^+^CD3^neg^CD19^neg^CD11b^+^MHCII^neg^) were 40.7% (SD±11.1) of the total CD45^+^ cells in circulation and dabrafenib treatment led to a drastic reduction in their absolute numbers (**Fig 8E**). Like the bone marrow, the majority of circulating MDSCs were Ly6C^int^Ly6G^hi^ PMN-lineage (83.9%, SD±6.6) while Ly6C^hi^Ly6G^neg^ m-MDSCs were a relatively minor fraction of the MDSC pool (4.0%, SD±1.9) (**Fig 8F** and **8G**). Dabrafenib treatment selectively decreased PMN-MDSC percentages in the blood (**Fig 8G**); which was reflected in the significant drop in absolute numbers of PMN-MDSCs per 1L of blood (**Fig 8H**). In the tumor microenvironment, dabrafenib treatment reduced MDSC numbers by 50% (**Fig 8I**). Like the blood, PMN-MDSCs were the majority MDSC population; however, there was an enrichment for m-MDSCs which were present at numbers comparable to PMN-MDSCs (∼3897 cell/mg tumor and 5455 cell/mg tumor for PMN- and m-MDSCs, respectively). In tumors from dabrafenib-exposed mice, the percentage of m-MDSCs increased by 50% with a corresponding decrease in PMN-MDSCs (**Fig 8J**). Absolute numbers of PMN-MDSCs per mg of tumor were reduced 7-fold by dabrafenib compared to controls while m-MDSCs showed a non-significant reduction in dabrafenib-treated animals (**Fig 8K**) Thus, the data suggests that dabrafenib reduces MDSC accumulation in the tumor microenvironment *in vivo* with a highly significant impact on PMN populations.

## Discussion

In this report, we describe an off-target effect of dabrafenib that impacts MDSC development by activation of the stress kinase GCN2, revealing developmental and transcriptional lineage relationships relevant for differentiation and function. Dabrafenib-induced GCN2 activity reduced ribosome association with transcripts and altered transcriptional programs preventing developmental transition from immature myeloid progenitors resulting in loss of PMN-MDSC differentiation and MDSC suppressive activity. Our findings are consistent with data from other groups showing cancer drives expansion of PMN-MDSCs from monocytic precursors, although we did not find that monocytic precursors expressed c-kit (CD117), suggesting in our model the monocyte PMN-MDSC precursors exist in a more mature state than the monocyte-like progenitors described by Mastio et al. (Mastio et al., 2019). Nevertheless, the data clearly shows an intimate relationship between PMN- and monocytic MDSCs, and highlights the functional balance between GCN2 activity, ISR signaling, and maturation of MDSC populations.

ISR signaling is critical for cells to balance demands of metabolic activity versus limitations of nutrient availability, protein translation and mitochondrial stress (McCormick et al., 2016). We have previously shown that loss of GCN2 in mature monocytic and PMN lineages alters MDSC transcriptional profiles with attenuation of immune suppressive function and increased expression of inflammatory cytokines (Halaby et al., 2019). This effect was indirect, resulting from transcriptional alteration of metabolic transcriptional programming reducing oxidative respiration (Halaby et al., 2019). In the current report we found that dabrafenib significantly increased general oxidative metabolism in MDSCs with a specific increase in oxidative phosphorylation transcriptional signatures in the same immature cell populations exhibiting inhibited developmental progression. This suggests that precise metabolic regulation is required to maintain inflammatory differentiation capacity, and dabrafenib-GCN2-driven hyperactivation of oxidative metabolism dysregulates inflammation-driven myelopoiesis.

The transcription regulator retinoblastoma protein (Rb) restricts PMN-MDSC development, controlling relative composition of monocytic versus PMN populations. In tumor-bearing conditions *Rb1* epigenetic silencing favors PMN-MDSC development (Youn et al., 2013) and its deletion promotes accumulation of monocytic precursors that can give rise to PMN-MDSCs (Mastio et al., 2019). We observed a significant reduction of the *Rb1* regulon in a key intermediate PMN cluster (cluster 9) after dabrafenib therapy. However, the *Rb1* regulon was not identified in the cluster 6 to cluster 9 trajectory branch suggesting differential Rb activity was not driving PMN population maturation. Moreover, cluster 9 showed the most significant dabrafenib-mediated loss of regulon activity indicating that transcriptional programming is highly impacted in this cell population.

There were relatively few regulons increased by dabrafenib with the most prominent being regulons that would be predicted to impact metabolism and cell identity (*Atf5*, *Mafg*, and *Zbtb7a*). In contrast, dabrafenib downregulated a number of regulons associated with *Jun* and *Fos* transcription factors, redox responses, metabolism, and cellular differentiation. While we observed a general reduction of ribosome association with mRNA, we did not observe a loss of relative polysome assembly, suggesting a decrease in translation but no specific inhibition of ribosome assembly. Thus, alterations observed are likely due to GCN2-induced transcriptional programs. The *Atf5* regulon was the only regulon increased in all dabrafenib-treated cell clusters suggesting strong, generalized induction. ATF5 is a key driver of the ISR transcriptional response, impacting mTOR function and oxidative metabolism in mammalian cells (Fiorese et al., 2016). Importantly, ATF5 also promotes cell survival, and its downregulation is required for astrocyte differentiation from neural progenitors (Angelastro et al., 2005). In addition, the amino acid starvation response is enriched in haematopoietic stem cells, promoting stem cell survival, with a rapid diminution as stem cells differentiate to common myeloid progenitors and finally to mature monocytic and PMN populations (van Galen et al., 2018). This correlated with reduced GCN2 expression in stem cells versus progenitor populations suggesting genetic control of GCN2 and eIF2χξ phosphorylation is directly associated with differentiation (van Galen et al., 2018). Thus, dysregulation of GCN2 and ATF5 activity would be expected to impact differentiation programs suggesting that this feature of dabrafenib may have prominent effects on MDSC differentiation.

Congenital loss of GCN2 does not impact basal immune cell composition in the periphery, and here we show GCN2 deficiency does not affect MDSC composition ± dabrafenib. This contrasts with the ISR kinase PERK, which is required for tumor-driven myelopoiesis and MDSC development in the spleen of mice (Liu et al., 2022). Complementation studies have suggested that GCN2 and PERK serve compensatory roles in sensing cellular stress (Hamanaka et al., 2005). Indeed, reactome analysis of MDSCs suggested dabrafenib induced a UPR and PERK transcriptional signature suggesting increased PERK activity; however, we previously reported that in MDSCs PERK does not drive compensatory nutrient starvation responses in the absence of GCN2 function, and likewise loss of GCN2 has no impact on UPR responses in the presence or absence of PERK (Halaby et al., 2019). This agrees with the data presented here showing that loss of GCN2 abrogated the effect of dabrafenib on ISR signaling and MDSC development. Thus, taken as a whole, the data suggests that the PERK and GCN2 branches of the ISR have distinct biologic roles in MDSC development and function. Our study shows a loss of suppressive activity when MDSCs develop in the presence of dabrafenib resulting from proliferative arrest and the loss of functionally competent MDSC populations. Likewise in this report we show that when dabrafenib is added to mature MDSCs there is no effect on the ability to suppress T cell responses. Coupled with the lack of impact on MDSC development, the data suggests that GCN2 function in the tissue microenvironment is an important factor regulating phenotype, but increased activity during development disrupts transcriptional and metabolic programs leading to attenuated development.

In this report we took advantage of off-target effects of dabrafenib to probe MDSC developmental relationships and function related to GCN2 activity. The data reveals novel insights regarding the relative balance of ISR signaling and MDSC development that may have therapeutic relevance. Dabrafenib therapy in BRAF-mut melanoma increases T cell infiltration and sensitizes the tumors to immune-checkpoint inhibition therapy (Frederick et al., 2013; Wilmott et al., 2012). However, dabrafenib does not affect human T cell or dendritic cell function *in vitro* suggesting alternative tumor-intrinsic or off-target effects are impacting anti-tumor immunity (Vella et al., 2014). Our data suggests immune-stimulating effects of dabrafenib therapy may be at least partially due to altered MDSC function, a prediction supported by the reduction of PMN-MDSCs in BRAF-mut tumor-bearing mice after dabrafenib treatment. Ultimately, a deeper understanding regarding how tumor-stroma interactions are impacted by dabrafenib in specific and off-target mechanisms will improve clinical management of the disease.

## Acknowledgments

Funding: This work was supported by NIH grants CA190449, AI105500, AR067763, the Medicine by Design/ Canada First Research Excellence Fund, The Terry Fox Research Institute, and grant PJT-162114 from the Canadian Institutes of Health Research (TLM).

## Author contributions

Conceptualization, T.L.M. and M.T.C.; Methodology, T.L.M. and M.T.C.; Investigation, M.T.C., R.Q., R.J., S.L., and N.N.; Writing – Original Draft, T.L.M. and M.T.C.; Funding Acquisition, T.L.M.; Resources, M.K.; Supervision, T.L.M.

## Figure titles and legends

**Supplemental Figure 1.** Gating strategy

**(A)** Gating strategy to analyse immune cell subpopulation in fresh bone marrow and MDSCs *in vitro* cultures.

**Supplemental Figure 2.** Top marker genes used for cluster identification.

**(A)** Normalized expression and per cluster percentage expression of top 10 marker genes for each cell cluster for the scRNA sequencing analysis on the integrated sample including MDSCs generated ± 1.5μM DAB.

**Supplemental Figure 3.** Selected regulon AUCell enrichment scores of cells in Ctrl or dabrafenib (DAB) conditions.

**(A)** Density plots representing the distribution of the AUCell score for dabrafenib-induced enriched regulons (*Atf5*, *Mafg* and *Zbtb7a*) and Ctrl-enriched regulons (*Mxd1*, *Tcf7l2* and *Fosl2*) in representative PMN (cluster 6 and 9) and monocyte clusters (cluster 12). Dashed line indicates geometric mean for Ctrl (black) or DAB (red). Exact q value and average Log2FC is indicated in each panel. No significance is indicated as “n.s”. Statistics show Wilcoxon test.

**(B)** Density plots representing the distribution of the AUCell score for dabrafenib-modulated regulons in monocytic cluster 12 (*Nfe2*, *Runx3* and *Tal1*). Dashed line indicates geometric mean for Ctrl (black) or DAB (red). Exact q value and average Log2FC is indicated in each panel. Statistics show Wilcoxon test.

**(C)** Heatmap of the Rb1 regulon during differentiation in the direction from node a to node b in Figure 6C, which spans from cluster 6 (immature granulocytes) to cluster 9 (intermediate granulocytes) in control cell. Scale represents z-score of the normalized average AUCell score per regulon.

**Supplemental Table A. Antibodies and reagents used for flow cytometry and FACSorting**

**Supplemental Table B. List of primers used in qPCR**

**Supplemental Table C. Antibodies used for western blot**

**Supplemental Table D. Data sets deposited on GEO repository used for annotation of scRNAseq clusters**

## Methods

### Mice

C57BL/6 mice were purchased from the breeding colony at the Princess Margaret Cancer Centre animal facility. *C57BL/6-Tg(TcraTcrb)1100Mjb/J* (OT-I), *B6.PL-Thy1a/CyJ* (Thy1.1) and B6.129S6-Eif2ak4tm1.2Dron/J (GCN2^-/-^) mice were purchased from The Jackson Laboratory. All mice were housed under specific pathogen–free conditions in accordance with the Canadian Institutional Animal Care and Use Committee guidelines. Protocols were approved by the Princess Margaret Cancer Centre Animal Care Committee. Female mice 7-10 weeks of age were used for all experiments.

### Tumor cell culture and tumor injection

Mouse melanoma YUMM1.7 tumor cells were obtained from ATCC. Cells were cultured in DMEM/F-12 (Gibco), supplemented with 10% FBS, 100 units/mL of penicillin, 100 µg/mL of streptomycin and 55 µM 2-Mercaptoethanol. Cells were maintained at 37°C in 5% CO_2-_humidified atmosphere. 3x10^5^ YUMM1.7 cells were injected subcutaneously (s.c.) into the right hind leg of each mouse.

### BRAFi formulation for *in vivo* treatments

Dabrafenib (30 mg/kg) (Cayman Chemical) or the equivalent volume of DMSO were dosed orally once a day as a suspension in 0.5% HPMC and 0.2% Tween80 in distilled water at pH 8.

### Generation of bone marrow-derived MDSC

Bone marrow was flushed out of the tibia and femur. Red cells were depleted with ACK lysing buffer. Cells were seeded at concentration 6x10^5^ cells/mL in RPMI 1640 medium supplemented with 10% FBS, 100 units/mL of penicillin, 100 µg/mL of streptomycin and 55 µM 2-Mercaptoethanol. GM-CSF (50 ng/mL) and IL-6 (50 ng/mL) were used to drive MDSC differentiation. Cells were maintained at 37°C in 5% CO_2-_humidified atmosphere for 4 days. Alternatively, myeloid progenitors were isolated with EasySep Mouse Hematopoietic Progenitor Cell Isolation Kit (StemCell) and stimulated in the conditions mentioned above. Increasing concentrations of dabrafenib or matching DMSO volume were added at the time of seeding.

### Suppression assays

OVA-specific CD8^+^ T cells were isolated from the spleens and lymph nodes of OTI/Thy1.1 mice by negative selection (Stemcell Technologies) and labeled with CFSE for proliferation tracking. CD11c^+^ cells were isolated from the spleens of C57BL/6 by positive selection (Stemcell Technologies) to serve as antigen presenting cells. Briefly, CD11c^+^ cells were pulsed with 1 µg/mL OVA_257-264 p_eptide for 1h at 37°C. After washing, cells were seeded at a ratio 1:10 CD11c^+^:CD8^+^ T cells in 96-well plates. MDSC were harvested and added to the cell mix at different ratios of MDSC:CD8^+^ T cells.

### Flow cytometry and cell sorting

Single-cell suspensions of cells were stained in a staining buffer (PBS, 2% FBS, 1mM EDTA) containing fixable viability Dye, Fc blocker and the appropriate antibody cocktail. Antibodies can be found in the extended Methods section. Samples were washed, fixed in 1% paraformaldehyde, washed again, and resuspended in staining buffer. Cells were acquired using a LSR Fortessa or LSR Fortessa X-20 analyzer (BD Biosciences) and data was analyzed using FlowJo software version 10.8.1 (Treestar). Alternatively, cells were FACSorted on FACSAria Fusion sorter (BD Biosciences) in tubes containing complete RPMI. See Supplemental Table A for antibody details.

### RNA isolation and quantitative rtPCR (qPCR)

RNA from lysates was purified using RNeasy Plus RNA purification kit (Qiagen) and retrotranscribed using qScript cDNA supermix (Quanta bio). cDNA was amplified using the PerfeCTa SYBR green Supermix (Quanta bio) on a CFXConnect real time PCR detection system (Biorad). Results were analyzed using the accompanying software according to manufactures instructions. See Supplemental Table B for primer list.

### Western blot

Fresh cells pellets were lysed in lysis buffer (1% Triton-X100, 50 mM Tris-HCl, 100 mM NaCl, pH 7.4) in presence of 1x protease and phosphatase inhibitors for 1h at 4°C while rotating. See Supplemental Table C for antibody details.

### Polysome Profiling

A total of 1x10^7^ cells per condition were used for polysome profiling as described previously (Ravishankar et al., 2015), with some modifications. In brief, cells were treated with 0.1 mg/ml cycloheximide (CHX) for 3 min at 37°C and washed twice with ice cold PBS/CHX. Cell pellets were then homogenized in 200 µL of lysis buffer (1% Triton X-100, 0.3 M, NaCl, 15 mM MgCl_2,_ 15 mM Tris (pH 7.4), 0.1 mg/mL CHX, 100 units RNAse-Inhibitors (Takara)) for 20 min at 4°C in a rotating mixer. The homogenate was centrifuged to remove nuclei, and supernatants were stored at −80°C until use. For each run, equal volumes of cell lysates were layered over a 10 mL linear sucrose gradient (20–50% sucrose in 15 mM, MgCl_2,_ 15 mM Tris (pH 7.4), 0.3 M NaCl) and centrifuged for 90 min at 39,000 rpm in an SW41-Ti rotor at 4°C. Absorbance at 254nm was measured continuously as a function of gradient depth in a BioRad Laboratories UV monitor. Overall translation efficiency was calculated as area under the region of the curve that represented mRNA attached to two or more ribosomes compared to the total area under the curve.

### Seahorse Assay

*In vitro* differentiated MDSCs were seeded in Agilent Seahorse XF96 microplates coated with 15 µg/mL of Cell-Tak (Corning) at a density of 1x10^5^ cells per well. Extracellular acidification rate (ECAR) and oxygen consumption rate (OCR) were measured after injections of 25 mM glucose, 1.5 µM oligomycin A, 1.5 µM FCCP, 1mM Sodium Pyruvate, 2.5 µM antimycin A, and 1.25 µM rotenone A optimizing a methodology previously described (Van den Bossche et al., 2015).

### Single-cell RNA-sequencing and analysis

MDSC were harvested and washed 2 times with PBS + 0.04% BSA. Then ∼1x10^4^ cells were captured with 10X Genomics Chromium Next GEM Single Cell 5’ Kit v2. Libraries were subsequently prepared and sequenced using a NovaSeq sequencer (Illumina).

The scRNA data was demultiplexed and converted to FASTQ file format with Illumina bcl2fastq. Initial quality control and alignment against mouse reference transcriptome Mm10-2020-A was performed on the resulting scRNA data using CellRanger (v7.0.0) count mode with SC5P-R2 chemistry. Briefly, Seurat (v4.3.0) was used to exclude any cells with >10% of the reads mapping to mitochondrial genes, cells that are considered as outliers (MAD>4) by expressing an excessive amount of genes/reads by scater (v1.22.0) (McCarthy et al., 2017), and cells/clusters considered as doublets as identified by the intersection of DoubletFinder (v2.0.3) (McGinnis et al., 2019) and scds (v1.9.1) (Bais and Kostka, 2020). We then performed normalization on the filtered count matrix and variance stabilization across 3000 genes, regressing out cell-cycle and mitochondrial genes, using SCTransform v2 (Choudhary and Satija, 2022). Samples were integrated across treatment conditions using FastMNN as implemented in SeuratWrappers (v0.3.1) and defined from the batchelor (v1.10.0) R package (Haghverdi et al., 2018). Principal component analysis (PCA) was performed, and the top 40 principal components were included in a Uniform manifold approximation and projection (UMAP) for dimensionality reduction, as well as for finding shared nearest neighbours and the Leiden Clustering algorithm with a resolution of 0.9. Differentially expressed genes between treatment conditions were calculated per cluster and cell-type using the Wilcoxon-test implemented in the FindMarkers function of Seurat using no fold-change or percent-expressed thresholds.

Cell type annotations were done by creating a reference atlas using 40 data sets that categorized features of developing and mature monocyte, PMN, and MDSC populations among others (Supplemental Table D). Each dataset was pre-processed using methods like those outlined in this paper, subSetted to have 1000 cells representing each dataset and integrated together using FastMNN. Cell-types were predicted per cell by estimating the shared nearest-neighbour between the query and reference datasets using ProjecTILs (v3.0.0) (Andreatta et al., 2021). These annotations were further refined by examining the differentially expressed genes across each cluster type.

Regulon analysis was performed using SCENIC (v1.2.4), pyScenic (v0.12.0), and their activity was scored using AUCell (v1.21.2) (Aibar et al., 2017). Genes were removed from this analysis if they were not expressed in at least 1% of all cells. Differential expression of regulons per cluster was performed using a Wilcoxon test between treatment conditions, corrected for multiple hypothesis testing using the Benjamini-Hochberg method, and regulons were deemed significant if they exceeded an |log2FC| > 0.01 and q < 0.01. Pathway analysis was performed on the MSigDB gene-sets Hallmark, C2:Reactome, and C5:GOBP, accessed through the msigdbr (v7.4.1) R package. Significant differentially expressed pathways were identified using SCPA v1.5.2 (Bibby et al., 2022) and their corresponding activity was estimated using AUCell.

Trajectory analysis was performed using monocle3 (v1.0.0) (Trapnell et al., 2014) by importing the UMAP and PCA coordinates from the pre-constructed Seurat object into the Single Cell Experiment object. Analysis was done according to the guide found at http://cole-trapnell-lab.github.io/monocle-release/monocle3. Selected branches were focused on by iterating through key start and end nodes, splitting the cells along that branch into treatment groups, and identifying variable genes along these paths with a q-value<0.05 and modularizing at a resolution of 0.001. To gauge the pathway and regulon activity across each branch, AUCell was used to score the activity of every cluster spanned by a given branch based on the average log-normalized expression of cells within that cluster and branch. Significant pathways and regulons were identified using an over-representation analysis as implemented in the clusterProfiler (v4.2.2) R package (Yu et al., 2012) of the gene-modules on these same gene-sets.

### Statistical analysis

Statistical parameters calculated with GraphPad Prism v9.5 are described in each figure legend. Error bars indicate the standard deviation (SD). IC50 values were calculated by nonlinear regression. Student’s t-tests or ANOVA were used for p-value calculations. Statistically significant differences (p<0.05) are indicated by the exact p-value in figures and legends.

### Data Availability Statement

Raw and processed single-cell RNAseq data is available from GEO under accession number GSE239496. Source code used to process the data and generate Figures 5 and 6 can be found at https://github.com/mcgahalab/ly6gc_mice.

