## Supplementary figures and images for "Dabrafenib alters MDSC differentiation and function by activation of GCN2"

### Supplemental Figures

#### Supplemental Figure 1

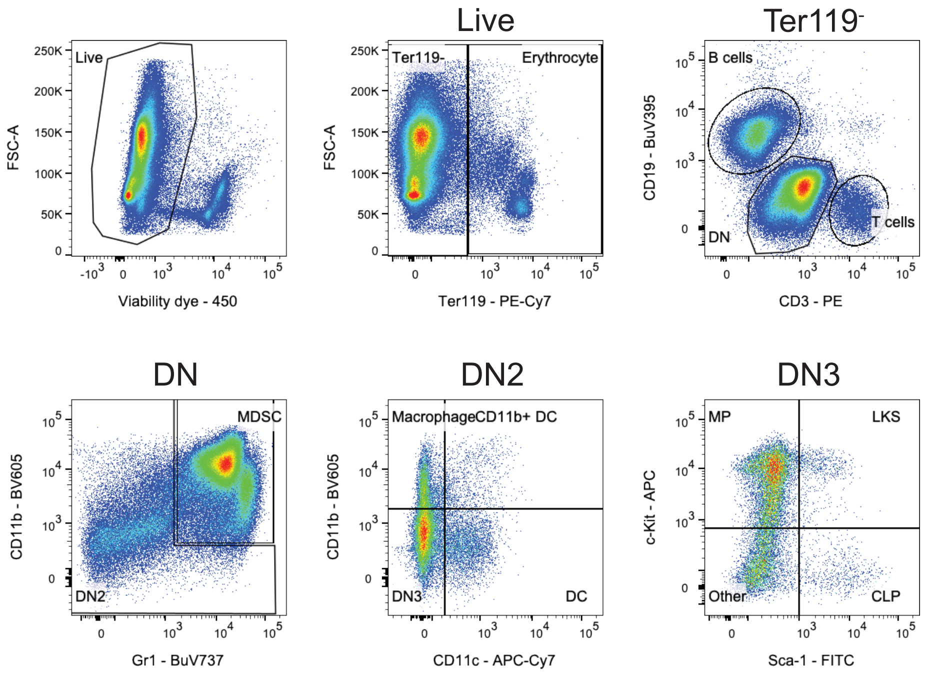

#### Supplemental Figure 2

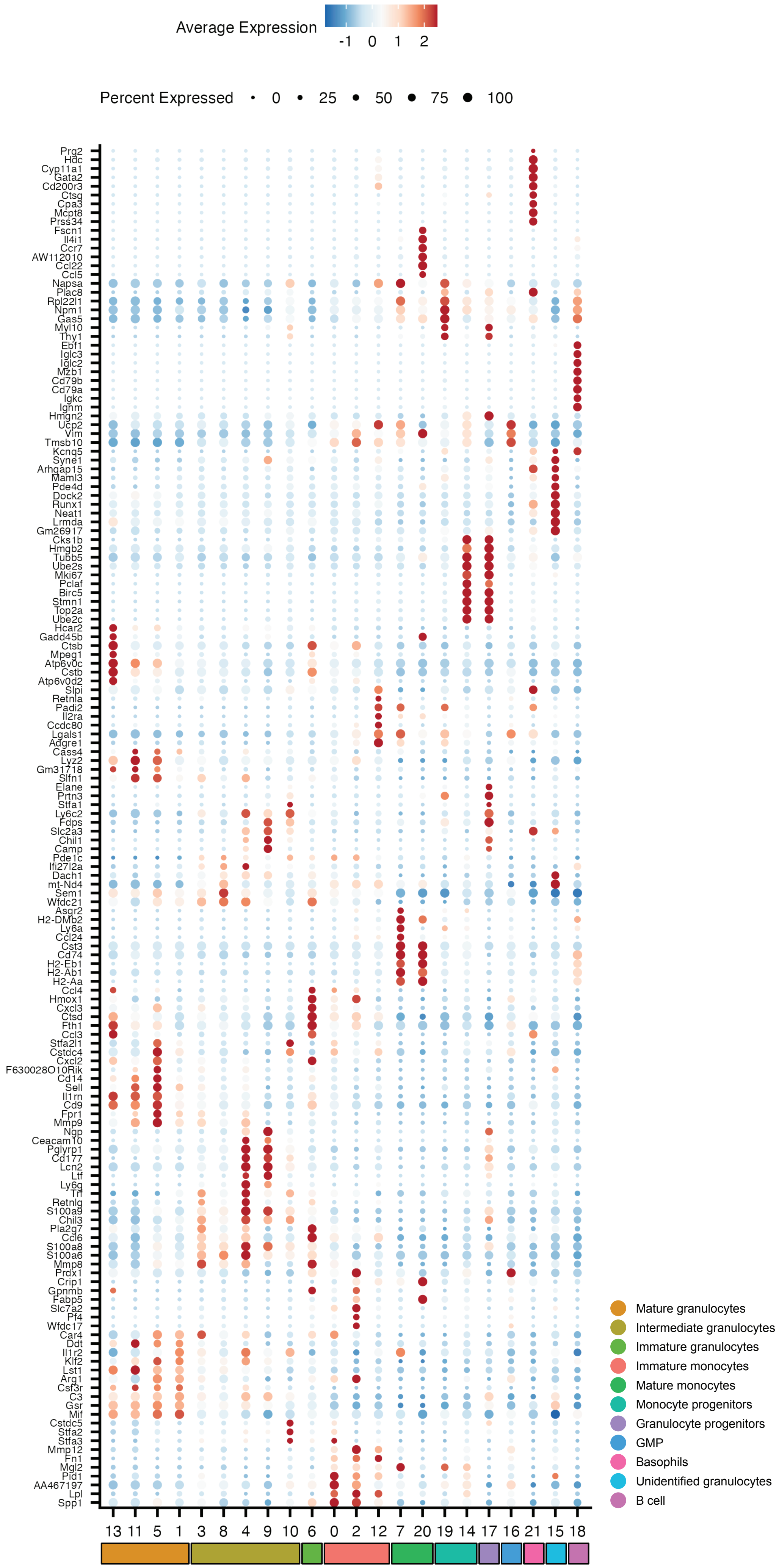

#### Supplemental Figure 3

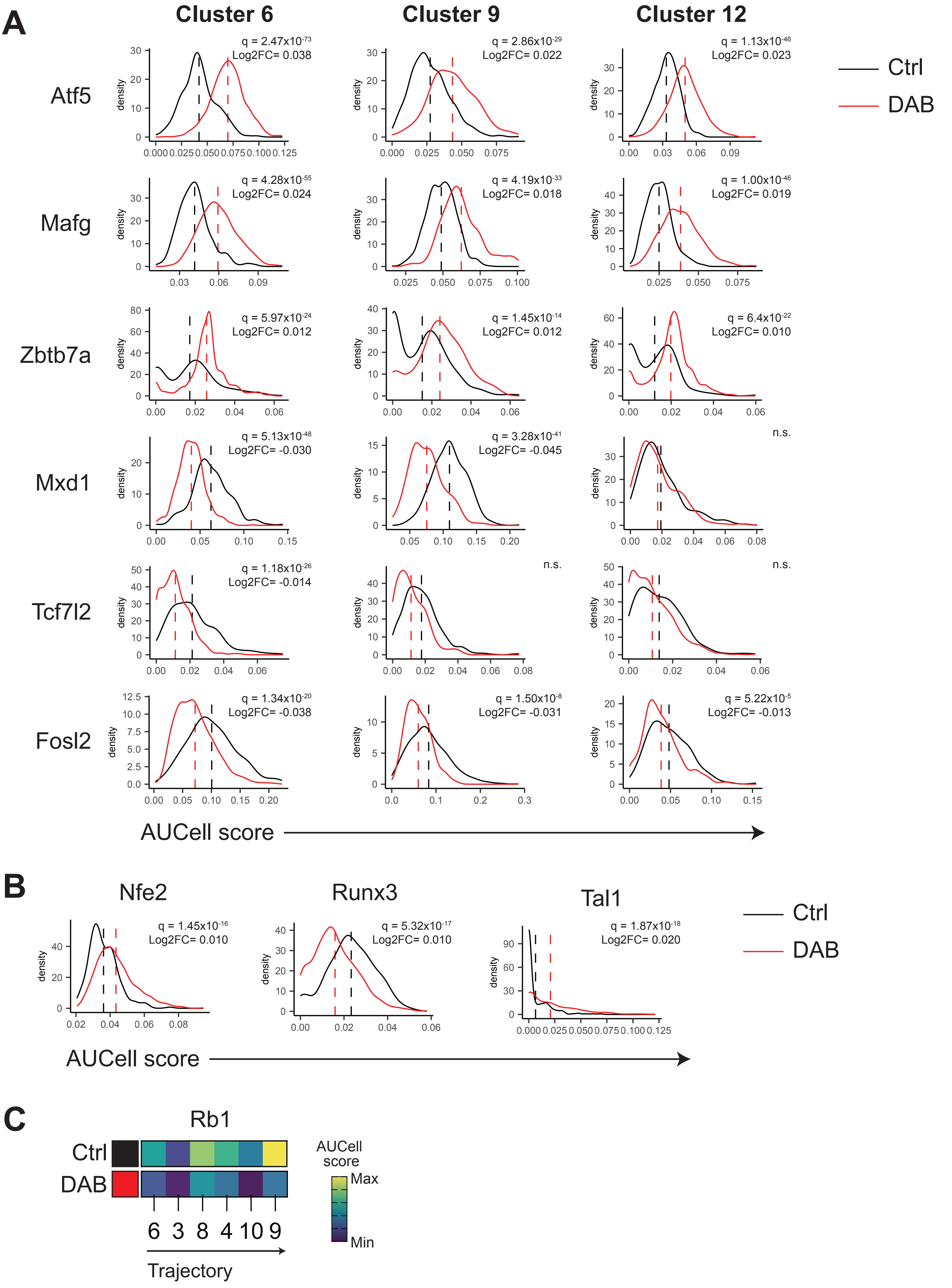
